# Massively parallel characterization and predictive modelling of neuronal regulatory variation

**DOI:** 10.64898/2026.07.16.738760

**Authors:** Kilian Salomon, Chengyu Deng, Pyaree Mohan Dash, Theofilos Chalkiadakis, Qinrui Li, Ziwei Chen, Nicholas F. Page, Mustafa Helal, Sebastian Röner, Anshul Kundaje, Claudia Langenberg, Maik Pietzner, Jay Shendure, Max Schubach, Nadav Ahituv, Martin Kircher

**Affiliations:** Computational Genome Biology, Exploratory Diagnostic Sciences, Berlin Institute of Health at Charité - Universitätsmedizin Berlin, Berlin, Germany; Department of Bioengineering and Therapeutic Sciences, University of California San Francisco, San Francisco, CA, USA; Institute for Human Genetics, University of California San Francisco, San Francisco, CA, USA; Department of Computer Science, School of Engineering, Stanford University; University of Luebeck, Institute of Human Genetics, University Hospital Schleswig-Holstein, Campus Luebeck, Germany; Computational Medicine, Medical and Health Data Sciences, Berlin Institute of Health at Charité - Universitätsmedizin Berlin, Berlin, Germany; Department of Genetics, School of Medicine, Stanford University; Precision Healthcare University Research Institute, Queen Mary University of London, London, UK; Health Data Modelling, Medical and Health Data Sciences, Berlin Institute of Health at Charité - Universitätsmedizin Berlin, Berlin, Germany; Department of Genome Sciences, University of Washington, Seattle, WA, USA; Seattle Hub for Synthetic Biology, Seattle, WA, USA; Brotman Baty Institute for Precision Medicine, Seattle, WA, USA; Howard Hughes Medical Institute, Seattle, WA, USA

## Abstract

Disease-associated variants reside frequently in noncoding cis-regulatory elements (CREs), yet their functional consequences remain poorly understood. We performed a large-scale lentiMPRA in human excitatory neurons, quantifying the impact of >46,000 naturally occurring variants across >27,000 candidate CREs near 524 disease-associated genes. These data improved regulatory variant effect predictions beyond state-of-the-art models. Significant allelic effects occurred at comparable rates across common, rare, and singleton variants, demonstrating that, within MPRA-measurable effects, population frequency carries limited information about per-variant regulatory impact. Variant effect detectability and magnitude were governed primarily by baseline activity of the enclosing regulatory element and local sequence context. Regulatory effects were distributed across numerous transcription factors rather than concentrated in master regulators, consistent with a combinatorial enhancer architecture. We establish a large-scale functional variant catalog and provide a complementary benchmark and resource for developing and evaluating models of noncoding regulatory variation.

## Introduction

Understanding the genetic basis of disease-associated variation is a key challenge in genomics ^1–3^. Completion of the human reference genome enabled large-scale functional genomics initiatives that systematically mapped chromatin accessibility, histone modifications, and transcriptional activity across diverse cell types ^4–6^. These efforts have cataloged millions of candidate cis-regulatory elements (cCREs), yet the precise mechanisms by which genomic sequence encodes cell-type-specific regulatory programs particularly within the vast non-coding genome remain incompletely understood ^7^.

Transcriptional regulation is governed by promoters and enhancers that integrate combinations of transcription factor binding sites (TFBSs) to control gene expression in specific cellular contexts ^8,9^. Resources such as ENCODE SCREEN provide epigenetic annotations of cCREs across tissues and cell types ^10,11^. At the same time, population-scale genotyping and sequencing has revealed that the majority of disease-associated variants reside in non-coding regions, frequently within putative regulatory elements ^3,12^. Experimental studies demonstrated that variants within cCREs can alter gene expression, underscoring their relevance for both common and rare disease phenotypes. However, only a small fraction of annotated cCREs and an even smaller fraction of human genetic variants have been functionally characterized in relevant cellular systems ^11^.

A central challenge is to determine which noncoding variants are functionally relevant and how their effects relate to population frequency ^13–16^. Population allele frequency is often used as a proxy for functional constraint, with rare variants expected to have larger effects under purifying selection ^17–20^ but allele frequency is shaped by demographic history and drift in addition to selection, and whether the assumed inverse frequency-effect relationship holds for regulatory variants in a defined cellular context remains poorly characterized. Computational predictions based on genomic location or TFBS are informative but incomplete, reflecting our limited understanding of regulatory sequence grammar ^14,21^. While CRISPR-based perturbation approaches enable in situ testing of selected loci ^22,23^, Massively Parallel Reporter Assays (MPRAs) allow high-throughput interrogation of thousands of regulatory elements and variants in parallel ^24^. Large-scale MPRA studies provide an opportunity to move beyond cataloging regulatory annotations toward quantifying regulatory activity and the functional impact of naturally occurring variants in a defined cellular context ^25^

Large-scale functional assays enable us to address several central questions. First, how well do current catalogs of cCREs define cell-type specific activity ^11,26^? Second, how do allelic regulatory effects in such elements relate to allele frequency, and to what extent can population frequency serve as a proxy for functional impact in regulatory DNA ^17,18^? Third, how well do current computational models capture these effects at single-nucleotide resolution ^14,21^? Furthermore, we examine how regulatory context influences variant effects, including the role of transcription factor binding, baseline regulatory activity, and mutation class. Together, these analyses aim to disentangle the contributions of sequence, context, and population genetic constraint to regulatory variation.

Here, we performed a large-scale lentivirus-based MPRA (lentiMPRA) in NGN2-derived human excitatory neurons to systematically interrogate regulatory architecture surrounding disease-associated genes. We tested over 27,000 cCREs located near more than 500 neural, cardiac, and clinically actionable genes and quantified their activity relative to scrambled controls. Within these elements, we assayed the functional impact of more than 46,000 single nucleotide variants (SNVs) from gnomAD v3.1.2, including over 22,000 common variants (allele frequency >1%) and over 24,000 rare and singleton variants. By integrating epigenetic annotations, transcription factor binding information, population genetic features, and sequence-based model predictions, we sought to quantify the sparsity and context dependence of regulatory activity and to evaluate how effectively current approaches capture functional non-coding variation in human neurons.

## Results

### MPRA of cCRE near disease-associated genes

We designed a large-scale lentiviral MPRA to interrogate regulatory elements and naturally occurring genetic variation surrounding disease-associated genes in human neurons (**Fig. 1**). Instead of starting with GWAS lead variants, the causal alleles of which are uncertain due to linkage disequilibrium, we based our design on disease-relevant genes. This approach allowed us to systematically investigate the cis-regulatory space around each locus and ensured that any detected allelic effect is proximal to a gene of established clinical or biological relevance. We selected a total of 525 genes across multiple disease-relevant categories, including 306 neurological genes (primarily from the SFARI database ^27^, supplemented with literature-curated genes associated with neuropsychiatric disorders such as schizophrenia ^28^, 164 cardiac genes (including congenital heart and coronary artery disease ^29–33),^ 48 clinically actionable genes (provided by the Center of Actionable Variant Effects, therefore abbreviated “CAVA”), and 25 randomly selected genes from GENCODE v42 (**Supplementary Table 1**). A small number of genes were shared across categories (10 neuro-cardiac, 3 neuro-CAVA, 3 cardiac-CAVA, and 1 overlapping all three categories), reflecting partially overlapping disease annotations.

**Figure 1:**
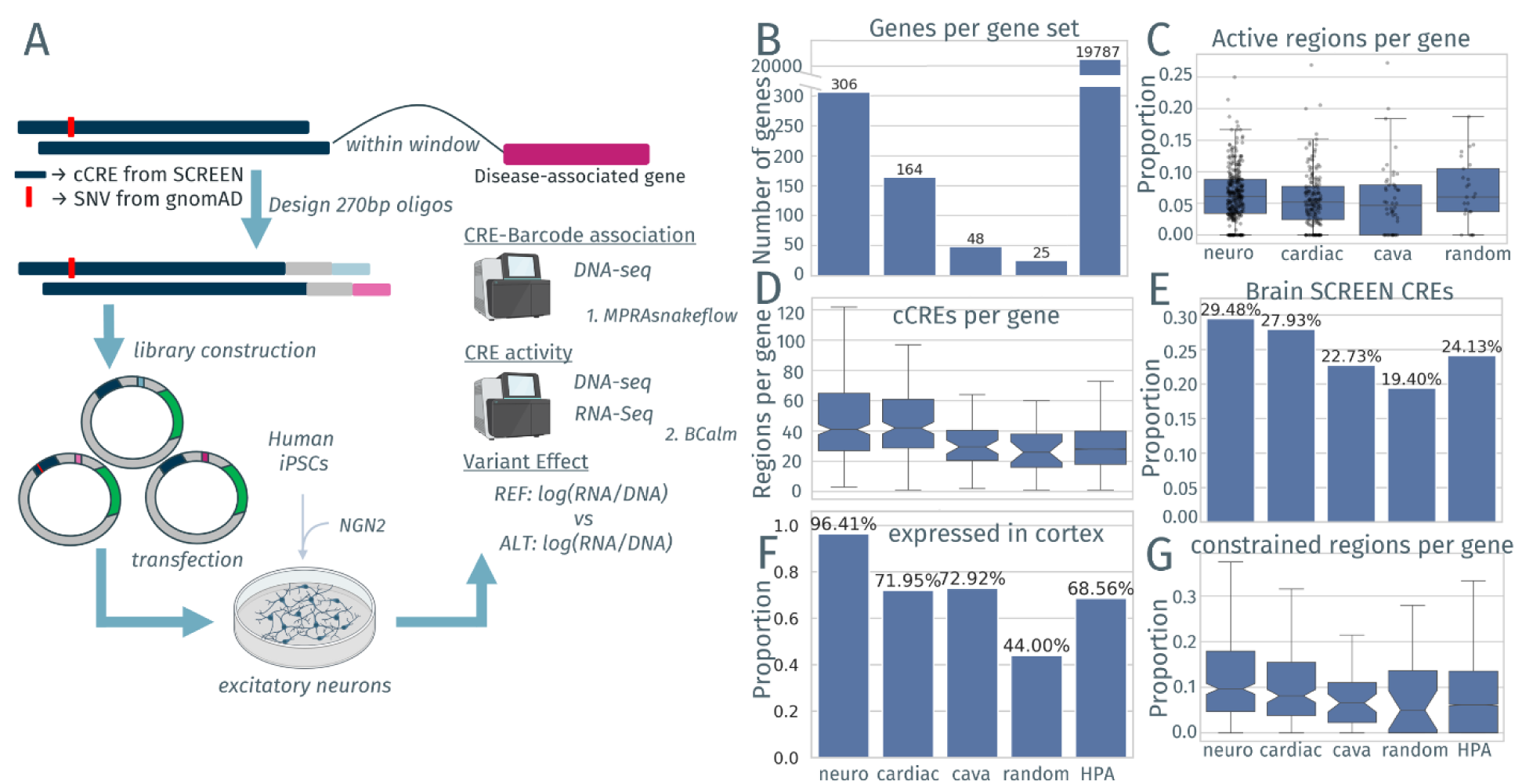
Design process and element overview. a, Outline of experimental and analysis approach including MPRA sequence design: Over 27,000 cCREs were designed, assigned to a random barcode in the 5’-UTR of a reporter gene and introduced into a lenti-vector. This library was sequenced to identify the barcode and sequence associations, and used to transfect the *NGN2*-induced neurons derived from WTC11. Targeted DNA-and RNA-seq and MPRAsnakeflow was used to quantify the barcode number on DNA-level (introductions into the genome) and RNA-level (observed transcripts of the reporter gene in the cell type). The ratio can be used to quantify the effect of the designed sequence on the transcription initiation of the reporter gene and the difference between the reference and alternative allele can be used to estimate variant effects (BCalm). **b**, Barplot of the number of genes in each of the gene sets, neuro (n=306), cardiac (n=164), cava (n=48), random (n=25), and HPA (n=19,787). **c**, Boxplot of the proportion of active cCREs per gene in each gene set. **d**, Boxplot of the number of cCREs per gene including all genes with data from human protein atlas (HPA; n=19,787) without showing outliers and plotting the confidence interval around the median. **e**, Barplot of the average proportion of elements considered as Brain SCREEN cCREs for each gene in the groups. **f**, Barplot of the proportion of expressed genes in cortex for each gene group. **g**, Barplot of the proportion of cCREs overlapping with constraint regions called Regions of contiguous constraint ^141^ per gene.

From these genes, we selected cCREs from the ENCODE SCREEN (v3) ^11^ that are located within 50 kilobase (kb) of their Transcription Start Sites (TSSs) (**Fig. 1a**). In total, 27,209 unique genomic regions were included in the design. Because cCREs are associated with genes on different strands, their sequences were designed on each strand respectively (27,482 including strand-specific duplicates). Within these regions, we designed 46,374 SNVs from gnomAD v3.1.2 spanning the whole allele-frequency spectrum. Because rare and singleton variants are substantially more abundant in the population than common variants, we applied an Enformer-based ^13^ prioritization strategy to enrich for potentially functional rare variants, selecting 70% variants with predicted activating effects, 15% with predicted repressive effects, and 15% randomly sampled variants for each gene set respectively. This design enriched for variants with predicted regulatory impact, introducing a bias for the rare variants. We therefore performed key analyses both on the full dataset and on the randomly sampled (unbiased) rare-variant subset to verify that conclusions were not driven by the prioritization (see below). We included multiple control sets to benchmark assay performance. Positive controls consisted of previously validated MPRA-active sequences and VISTA enhancers with neural activity (n = 166) ^34–36^. Negative controls included scrambled sequences (n = 500) and previously characterized low-activity elements (n = 451), as well as additional Impact of Genomic Variation on Function (IGVF) consortium defined controls (see Methods).

Sequences were synthesized as 270 bp oligos, cloned into lentiviral reporter constructs, and transduced into NGN2-derived human excitatory neurons differentiated from WTC11 iPSCs in three biological replicates (Methods). Regulatory activity was quantified from RNA-to-DNA barcode ratios. Of the 76,982 designed oligos, 67,904 (88%) passed quality control (see Methods), and sequences supported by at least 10 barcodes were retained. Replicates showed high concordance (Pearson r > 0.86), confirming assay robustness. After quality control and filtering, 24,358 unique elements (24,510 including elements tested in both strand orientations) yielded measurable activity. Most elements were associated with neurological genes (n = 16,151), followed by cardiac (n = 7,546), clinically actionable genes (CAVA, n = 1,610), and random genes (n = 717), with additional elements linked to overlapping gene categories. Of the 46,374 designed SNVs, 38,684 unique variants (38,968 including variants tested in both strand orientations) were successfully assayed. These included 10,309 singleton variants (allele count = 1), 10,251 rare variants (AF ≤1%), 8,077 common variants (AF 1-5%), and 10,047 very common variants (AF >5%).

To further characterize the composition of the dataset across gene groups, we examined several properties of the selected genes and associated cCREs (**Fig. 1b-g**). Gene categories differed substantially in size, ranging from 25 randomly selected genes to 306 neurological disease genes (**Fig. 1b**). Despite these differences in category size, both the number of cCREs tested per gene and the proportion of active elements per gene were broadly comparable across groups (**Fig. 1c, d**), indicating that differences in downstream analyses are unlikely to be driven by systematic differences in region representation or assay sensitivity. As expected from their tissue-specific selection, neurological genes showed the highest proportion of elements overlapping brain cCRE annotations (29%), yet substantial overlap was also observed in other gene sets, with an average of 24% across genes from the Human Protein Atlas (HPA), suggesting that brain-active regulatory elements are not exclusively concentrated in the neurological gene set (**Fig. 1e**). Consistent with their selection criteria, the neurological gene set showed the highest fraction of genes expressed in the cortex (96%), while the random set showed the lowest (44%), and HPA genes occupied an intermediate level (67%), confirming that the gene sets capture a meaningful gradient of brain relevance (**Fig. 1f**). Finally, evolutionary constraint scores of the associated cCREs were broadly similar across all gene groups, with no group showing a substantially elevated or depleted proportion of constrained regions (**Fig. 1g**), suggesting that the gene sets are subject to comparable levels of evolutionary constraint and that this variable is unlikely to confound cross-group comparisons in downstream analyses. Overall, this resulted in a dataset comprising 24,358 cCREs and 38,684 variants with quantitative activity measurements, spanning a broad range of allele frequencies and genomic contexts.

### Regulatory activity of cCREs in NGN2-derived neurons

We observed that the tested cCREs were on average only slightly more active compared to a scrambled control, while our controls selected for high and low activity separated well (**Fig. 2a**). Focusing only on reference allele elements of designed cCREs, we identified 2.1% (510/24,510) with significantly increased and 4.8% (1,178/24,510) with significantly decreased activity compared to scrambled controls (FDR < 0.05, BCalm ^37^). In the following, we refer to cCREs with significant activity differences relative to scrambled controls as differentially significant CREs (dCREs).

**Figure 2:**
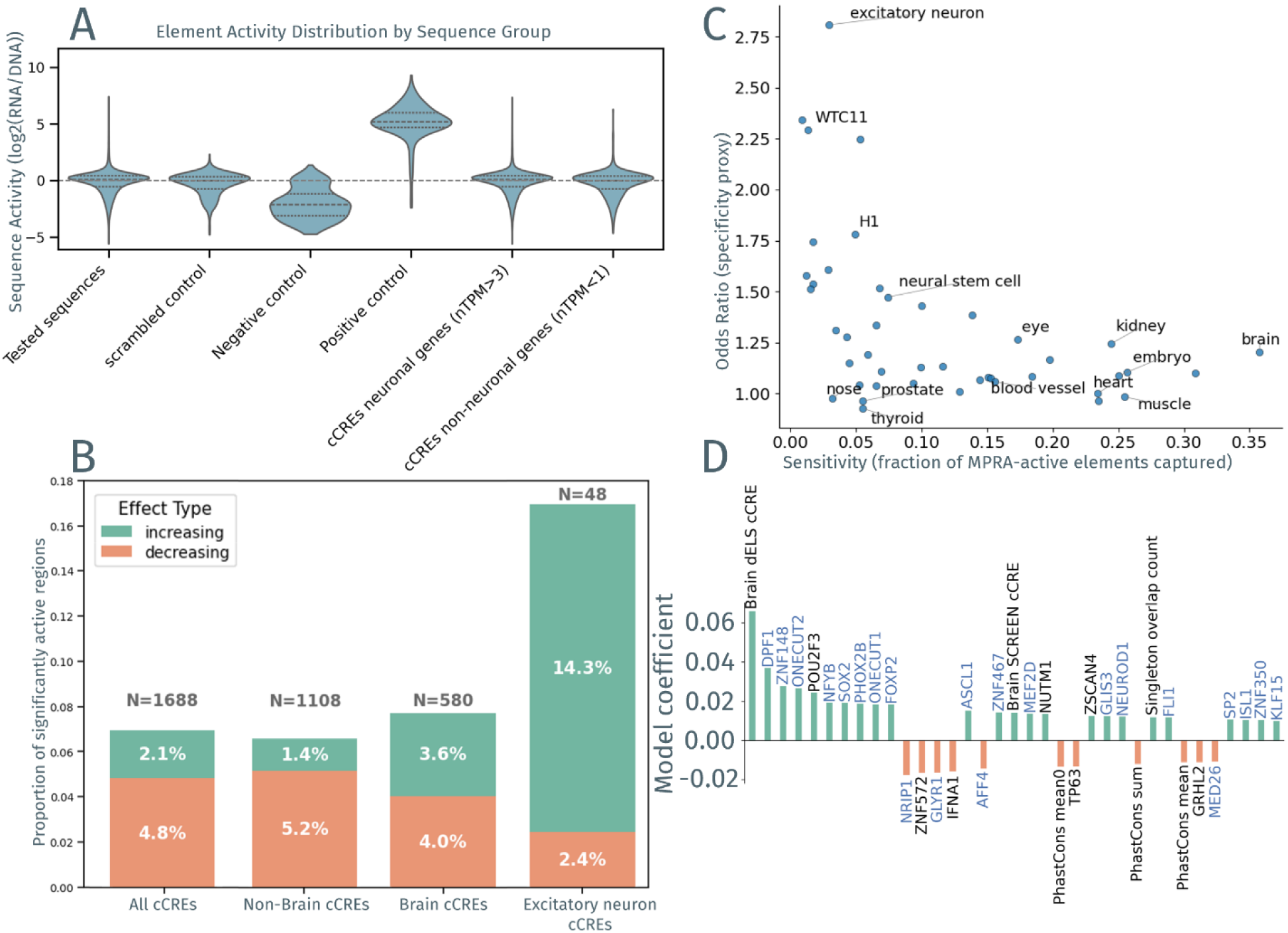
Element activity investigation. **a**, Activity distribution of the tested cCREs (n=65,308), a scrambled sequence set (n=474), negative (n=269) and positive (n=166) controls based on previous MPRAs as well as regions around neuronal genes (>3 nTPM in HPA brain tissues; n=22,480) and genes not expressed in brain related tissues (<1nTPM in HPA brain tissues; n=960). **b**, Proportion of significantly active sequences (compared to scrambled; n=1688) of sequences with (n=580) and without (n=1108) additional epigenetic marks in brain as well as excitatory neurons (n=48), according to ENCODE SCREEN (v4). **c**, Scatterplot of SCREEN cell-type performance based on the assessment of sensitivity, defined as the fraction of MPRA-active cCREs captured by a given annotation, and specificity, defined as the enrichment of MPRA-active cCREs within that annotation relative to all tested cCREs for the 38 major cell types and tissues SCREEN is organized in and additional cell-type specific cCREs considered to be close to the cell type tested in, neural stem, neural progenitor, WTC11, H1, excitatory neuron. **d**, Top 35 feature coefficients predicting element activity sorted by absolute value in a regularized linear model (Elastic Net) trained on TFBS occurrences and region-based features. TFs with ≥5 nTPM in at least one brain region according to HPA were highlighted with blue.

To assess whether regulatory activity depends on the expression status of the associated gene, we stratified cCREs based on gene expression in the human cortex (Human Protein Atlas). A total of 92% (22,480/24,510) cCREs were associated with genes expressed in neurons (nTPM >3), while 4% (960/24,510) cCREs were linked to genes with low or no neuronal expression (nTPM <1). As expected, activity distributions were shifted toward higher activity in the cCREs associated to neuronal expressed genes (median shift 0.07, p-value 0.002). With both sets having an interquartile range of 0.95 and 1.14 we note that this shift is small (**Fig. 2a**), indicating that MPRA-measured regulatory activity is not restricted to cCREs near actively expressed genes, but instead reflects intrinsic sequence-driven regulatory potential. To contextualize activity in NGN2-derived neurons, we annotated elements with the newer ENCODE SCREEN v4 release ^11^. Of all tested sequences, 30.9% (7,583/24,510) overlapped cCREs from brain tissues and brain-derived cell types (see Methods). Among the significant elements, 34.4% (580/1,688) were brain cCREs, indicating that elements annotated as open chromatin in brain by SCREEN are slightly more likely (OR = 1.18, Fisher’s exact test, p = 0.001) to display regulatory activity in NGN2 neurons (**Fig. 2b**).

The directionality of regulatory effects differed markedly between elements annotated as brain cCREs and those lacking cCRE annotations in brain according to SCREEN. Brain cCREs showed a near-equal distribution of activity-increasing and activity-decreasing effects (3.6% versus 4.0%), whereas non-brain cCREs were strongly enriched for activity-reducing elements (OR = 3.35, Fisher’s exact test, p ≈ 5.4e-28). These findings indicate that elements accessible in neuronal or brain tissues encode both activating and repressive regulatory functions, while elements annotated outside the brain predominantly constrain transcriptional output when tested in NGN2-derived neurons.

To contextualize the substantial fraction of dCREs lacking brain annotation (65.6%), we examined their distribution across SCREEN tissue categories. Although brain cCREs represented the single most frequent annotation among dCREs (n = 580), substantial numbers of active elements were also annotated in blood (n = 501), skin (n = 417), muscle (n = 413), embryo (n = 406), kidney (n = 397), and lung (n = 381), indicating that regulatory activity in NGN2-derived neurons is not restricted to elements annotated in neural tissues (**Extended Data Fig. 1**). This observation may partly reflect incomplete sampling of neuronal cell states in current annotation resources ^11^. Alternatively, it could also relate to sequences being inactivated by chromatin context, thus only showing activity in the MPRA experiment and enabling readouts of variant and element activity outside the endogenous cell profiles ^38^.

Benchmarking epigenomic annotations revealed a precision-recall trade-off: brain cCREs captured 34% of active elements with modest enrichment (OR = 1.18, Fisher’s exact test, p = 0.001), whereas excitatory neuron-specific annotations were highly enriched (OR = 2.8, Fisher’s exact test, p ≈ 1.5e-7) but covered fewer than 3% of active elements (Fig. 2c), underscoring the incomplete representation of neuronal regulatory states in current epigenomic catalogs.

To explore mechanistic underpinnings, we tested whether known TFs derived from ReMap2022 ^39^ were differentially associated with brain-annotated versus non-brain active elements, adjusting for GC content and total TF density (see Methods). Brain-annotated active elements were strongly enriched for binding sites associated with neurodevelopmental transcriptional programs, including the neuronal lineage regulators SOX4 ^40^, SOX21 ^41^, ASCL1 ^42^, NEUROG2 ^43^, PAX6 ^44^, POU3F2 ^42^, RORβ ^45^, and EBF3 ^46^, as well as chromatin-associated regulators such as CHD7 ^47^ and SMARCA2 ^48^ (OR = 3.45-9.71, ridge logistic regression, bootstrap FDR < 0.05). These factors are broadly implicated in neuronal specification, differentiation, and chromatin remodeling during neural development, consistent with the regulatory identity of NGN2-derived neurons. In contrast, non-brain annotated elements were enriched (i.e., brain-annotated elements were depleted, adjusted OR < 1) for factors associated with chromatin remodeling and corepressor-linked regulatory programs, including the NuRD complex component GATAD2B, which mediate histone deacetylation and chromatin compaction ^49,50^, as well as corepressor-associated regulators such as TBL1XR1/TBL1X, components of the NCoR/SMRT complex implicated in transcriptional silencing during neural development ^51,52^, and the nuclear receptor NR2F1, a regulator of cortical identity with context-dependent repressive activity ^53^ (OR=0.22-0.34, ridge logistic regression, bootstrap FDR < 0.05) (**Supplementary Table 2**). Together, these results suggest that regulatory sequences lacking brain-specific chromatin accessibility are preferentially associated with chromatin-modifying and corepressor complexes that may limit transcriptional activation in neuronal contexts.

### Analysis of CRE sequence features

To identify sequence features contributing to transcriptional activity, we annotated tested cCREs with known TFBS from ReMap2022 ^39^ and used the TFBS occurrences per cCRE together with genomic context-based annotations to predict element activities with a regularized linear model (Elastic Net; see Methods). Because a large proportion of elements showed activity values near zero, the model explained only a modest fraction of the total MPRA activity variance (test R² = 0.03 across all elements; 0.12 on a balanced active/inactive set; see Methods and **Extended Data Fig. 2**), with brain cCRE annotation and the expression of brain-enriched TFs among the strongest positive predictors (**Fig. 2d**). We also note that such a simple linear model is restricted to additive and context-averaged interaction contributions and does not explicitly resolve TF-TF interactions, motif positioning, or strand orientation. Among the 35 largest (absolute) coefficients, most (n = 24) contributed positively to activity (**Fig. 2d**). This includes whether the region was overlapping a brain CRE and several TFs highly expressed in brain (≥5 nTPM; see Methods). Some notable examples are DPF1, a chromatin remodeler of post-mitotic neurons that is specifically expressed in the brain ^54^, ONECUT2, a TF that when overexpressed in fibroblasts induces neuron-like morphology and expression of neuronal genes ^55^, or NRIP1 (with the highest negative coefficient), a ligand-dependent co-repressor of nuclear receptors ^56^ that is expressed in the brain, enriched in cortex and hippocampus, and required for hippocampus-dependent learning and memory in mice ^57,58^. PhastCons sequence conservation ^59^ showed the largest negative coefficient, but this sign reflected a dataset composition effect rather than a suppressive relationship with regulatory activity. Both activity-increasing and activity-decreasing dCREs were significantly more constrained than non-significant elements (median phastCons_mean 0.076 and 0.084 vs. 0.047; Mann-Whitney U, BH-corrected p < 10-7 for both), and the overall fraction of significant elements nearly doubled across phastCons quartiles (5.1% in the least constrained to 10.1% in the most constrained quartile; χ² = 134, p < 10^-26^). The negative direction of the elastic net coefficient arose because activity-decreasing dCREs outnumber activity-increasing ones, biasing the signed regression toward lower predicted activity in highly constrained regions. A full list of non-zero coefficients is available in **Supplementary Table 3**.

Limiting our analysis to dCREs versus elements not different to scrambled sequences and performing an enrichment analysis (Fisher’s exact test; BH FDR < 0.05), we identified 81 TFs with differential binding site representation (odds ratio > 2 or < 0.8, 54 or 27). Among these, 16 enriched TFs were found among the top 30 TFs ranked by absolute Elastic Net coefficients (**Fig. 2d**). Overall, 37% (30/81) showed non-zero model coefficients, with 36% (20/81) positive and 6% (10/81) negative coefficients (**Extended Data Fig. 3**). We conclude that while there is some concordance between motif enrichment and model-derived feature importance for predicting the full activity range, certain TFs specifically contribute to the activity of the most active or repressive elements (**Supplementary Table 4**). Among the 30 TFs which showed enrichment in dCREs, we found the aforementioned chromatin remodeler DPF1, ZSCAN4, a known transiently expressed TF in stem cells with binding sites conserved in microsatellites within mouse and human ^60^, and PHOX2B, a transcription factor involved in neuronal lineage specification and differentiation of autonomic neurons ^61^.

### Allelic regulatory effects across common and rare variants

To quantify how the frequency of naturally occurring variants relates to regulatory activity and to assess whether allele frequency is linked to functional impact, we analyzed allelic effects across the full set of tested variants. We tested our comprehensive sampling of gnomAD variants with AF >1% together with the tested subset of rare and singleton variants within each cCRE. Of the 46,374 designed variants, 38,968 (84%) had sufficient coverage for testing allelic effects. Comparing activity between alleles, we identified 1,018 variants (2.6%) with significant allelic effects (FDR < 0.1), including 605 variants with increased activity and 413 with decreased activity. Effect sizes are defined by the fold change between alternative and reference allele activity (log2FC) ranged from -1.5 to 1.65, corresponding to an approximately 0.35-fold decrease to a 3.14-fold increase in activity. Unless otherwise stated, allelic analyses were performed on all variant readouts passing quality filters.

Because cCREs associated with genes on different strands were designed in both orientations, 271 variants were represented twice in opposite orientations, and an additional 13 variants were tested twice within overlapping cCREs on the same strand. Across these 284 duplicated variants, 6 showed a significant allelic effect (FDR < 0.1) in at least one instance. The 13 same-strand duplicates included 2 variants reaching significance in both cCRE contexts, with concordant effect direction in both cases. Among the 271 strand-pair variants, 4 reached significance with concordant effect direction in 3 of 4 cases. For the one case that did not show concordant effect, both DNA and RNA of the non-significant variant were supported by only a fraction of barcodes compared to the barcode number of the significant variant (24/115 and 41/74), suggesting lower detection power. This indicated that neither strand representation nor cCRE context introduced a major bias in downstream analyses.

The dataset captured ∼83% of common gnomAD SNVs (AF >1%) in the targeted cCREs and 1.7% of rare variants, providing near-complete common variant coverage alongside a prioritized rare variant subset. Across allele-frequency groups, 2.1% of very common variants (AF >5%), 2.6% of common variants (AF >1%), 2.8% of rare variants (AF ≤ 1%), and 3.0% of singletons (AC = 1) showed significant allelic effects. We note that rare and singleton variants were prioritized using Enformer. When restricting analyses to only randomly sampled variants, the proportion of significant effects in rare and singleton variants decreased (1.9% and 2.3%, respectively). Because the unbiased subset contained far fewer rare variants (n = 3114), this comparison had substantially reduced power and did not allow a firm conclusion on whether rare variants are more likely to show significant allelic effects. Within the AF >1% portion of the design, common variants (AF 1-5%) showed a higher proportion of significant variants than very common variants (AF >5%) (odds ratio 1.23; Fisher’s exact p < 0.043; **Fig. 3a**), supporting a modest depletion of significant variant effects with higher allele frequency. Most importantly, the dataset showed that significant effects are detected across all allele-frequency ranges and at similar rates (**Fig. 3a-b**).

**Figure 3:**
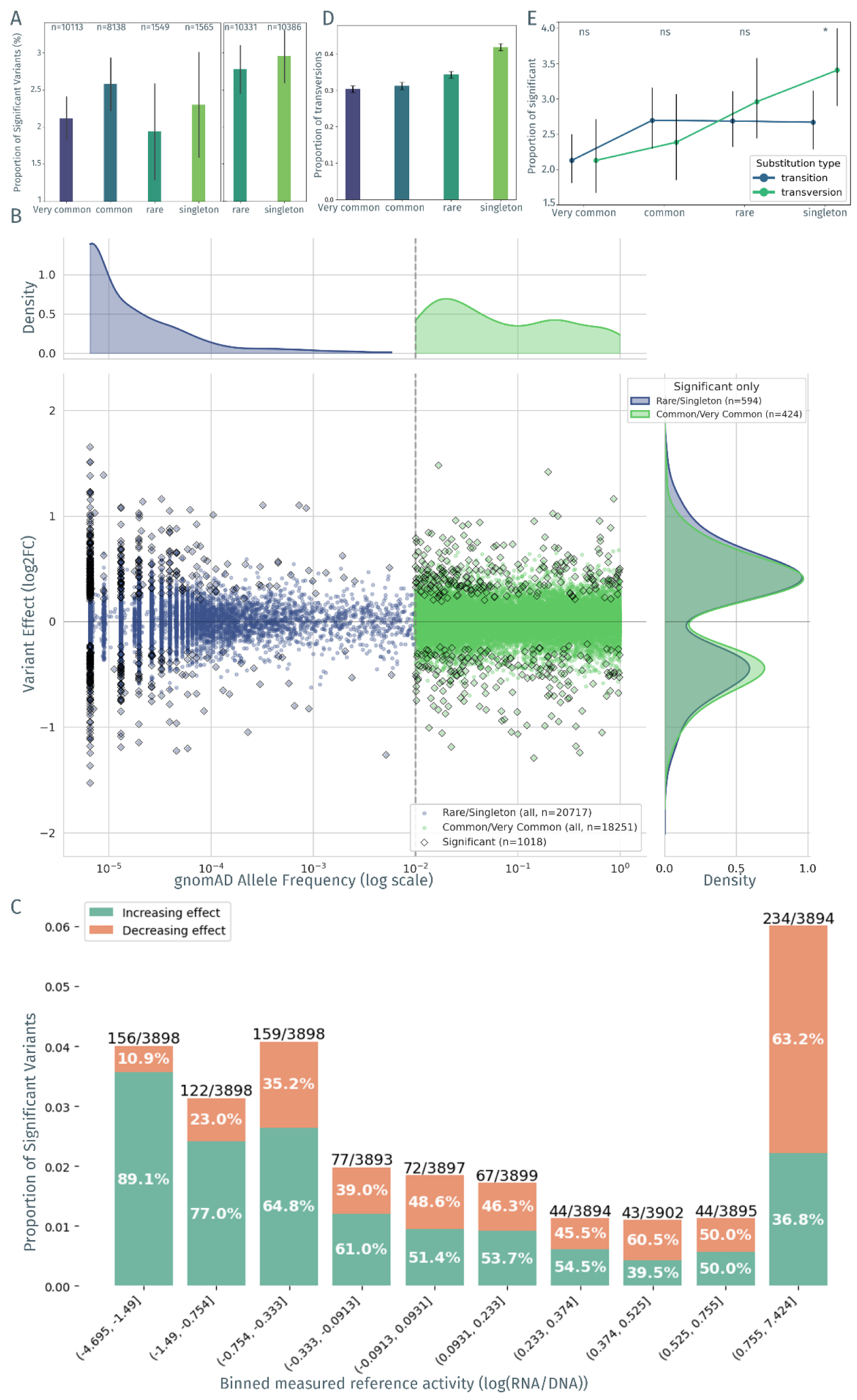
Stratified variant proportions and effect relationships. **a**, Proportion of significant variant effects per allele frequency (AF) group with AF >5% (“very common”, n=10,113), AF >1% (“common”, n=8,138), AF <= 1% (“rare variants”, n=10,331) and “singletons” (i.e. allele count (AC) of 1, n=10,386). Most rare variants were prioritized using Enformer during the MPRA design and only the set of randomly sampled variants is shown on the left (“rare”, n=1,549, “singleton”, n=1,565), while on the right all tested variants within the allele frequency group are included. AF and AC are derived from gnomAD (v3.1.2). **b**, Scatterplot of the relationship between variant effect direction and log-scaled AF derived from gnomAD (v3.1.2) for the full variant dataset (n=38,968) with rare and singleton variants in blue (n=20,717), (very) common variants in green (n=18,251), and significant effects represented as black diamonds (n=1,018). On top, the allele frequency distribution (left) and variant effect distributions (right) is shown for significant allelic effects (FDR<0.1) only. **c**, Proportion of increasing and decreasing variant effect with respect to the measured reference sequence activity, each decile activity bin is represented by ∼3900 variants. **d**, Proportion of transversion with respect to the allele frequency groups. **e**, Proportion of significant allelic effects between transition and transversion within allele frequency groups. Significance is presented by * (<0.05) or non-significant (ns).

Next, we asked whether allele frequency is related to the magnitude or direction of allelic effects. We modeled absolute effect size as a linear function of log_10_(AF), fitting rare (AF ≤ 1%) and common (AF > 1%) variants separately because the spectrum is sampled at very different densities in the two regimes. No significant relationship was observed for rare variants in either the full dataset (β = 0.0014, p = 0.36) or the randomly sampled subset (β = 0.0044, p = 0.21; slope-difference z-test, p = 0.43), and only a weak non-significant positive trend in common variants (β = 0.0025, p = 0.068). A single fit across the full AF spectrum showed a weak negative trend in the prioritized dataset that disappeared in the random subset (β = −0.0010, p = 0.001 vs β = 0.0006, p = 0.28; slope-difference p = 0.010), consistent with prioritization rather than population constraint (**Extended Data Fig. 4**). For signed allele effects, no consistent relationship was observed. Among significant variants, rare and singleton variants showed a somewhat higher proportion of activating effects than common and very common variants (61.6% versus 56.4%), but this difference was not statistically significant (χ² = 2.61, p = 0.106) (**Fig. 3b**). Extending this analysis across all allele frequency categories, we observed a significant monotonic decrease in the proportion of activating variants from rare to common variants (singleton: 62.9%, rare: 60.3%, common: 60.0%, very common: 52.8%; Cochran-Armitage trend test; Z = -2.16, p = 0.031), again indicating a small effect of allele frequency on the regulatory effect.

We observed that the detection of allelic effects depended on the transcriptional activity of the reference allele. Variants located in low-activity cCREs showed a higher proportion of activity-increasing effects, whereas variants in highly active cCREs more often showed activity-decreasing effects (**Fig. 3c**). While this asymmetry was partly explained by the limited dynamic range near baseline activity in MPRA constructs, where decreases are inherently harder to detect, it also reflects a biologically meaningful constraint, as highly active elements may be closer to a functional ceiling and therefore have limited capacity for further activation. In line with this, previous studies have preferentially focused on variants within active elements, often referred to as expression-modulating variants (emVars) ^62^, where regulatory effects are more readily interpretable, although our results indicated that substantial and statistically robust effects were also present outside this subset (**Fig. 3c**).

Further, we examined mutational classes, i.e. the classification of SNVs into transitions and transversions, reflecting substitutions within purines or pyrimidines, or between them. In our set of designed gnomAD variants, the proportion of transversions decreased steadily with increasing allele frequency (**Fig. 3d**), consistent with transversions being more strongly constrained, leading to their depletion at higher allele frequencies ^63–66^, even when removing CpGs (**Extended Data Fig. 5**). When comparing proportions of significant allelic effects between transitions and transversions among all tested variants, we did not see significant shifts, except for singleton variants where transversions exhibited a nominal significantly higher proportion than transitions (odds ratio 1.29; two-sided Fisher’s exact p < 0.035, **Fig. 3e**) suggesting a modest enrichment of functional effects among more strongly constrained mutation classes at low allele frequencies.

Overall, these analyses indicate that allelic regulatory effects are observed across the full allele-frequency spectrum, with only weak dependence on allele frequency for both effect size and direction. Instead, the detectability and characteristics of variant effects are more strongly influenced by local regulatory context, including baseline activity of the surrounding element. Together, these findings suggest that population frequency provides limited information about regulatory impact, while sequence context and intrinsic element activity play a central role in shaping measurable allelic effects in our MPRA study.

### Regulatory context and prioritization of variant effects

To characterize the regulatory context of significant allelic effects, we annotated all tested variants with regulatory element annotations (SCREEN cCREs) ^10^, enhancer-gene (E2G) links ^67^, fine-mapped eQTLs ^15^, GWAS loci ^68^, predicted TFBS (FIMO+ HOCOMOCO v13), and experimentally supported TFBS (ReMap2022) ^39^.

Enrichment across annotation classes (e.g. SCREEN cell-type annotation, presence/absence of E2G link, overlap/no overlap with eQTL/GWAS/predicted TFBS) measured as odds ratio was generally below 1.5 (**Extended Data Fig. 6**). Variants overlapping SCREEN brain cCREs were significantly enriched among significant variant effects in the MPRA (40.5% vs. 34.4%; OR = 1.31, FDR < 0.001), as were cCREs annotated for H1-cells (OR = 1.42, FDR = 0.04; **Extended Data Fig. 6**). In contrast, E2G links, GWAS loci, brain eQTLs, and ReMap2022 TFBSs did not show an enrichment.

Nearly all significant variants (94.8%) overlapped at least one predicted TF motif with local bit score filtering, a slightly lower overlap was observed for all tested variants (93.3%). Given the known false-positive rate of PWM-based motif scanning ^69,70^, the similarity of motifs among related TFs ^71,72^, and incomplete motif coverage across the TF repertoire ^73,74^, this near-saturating overlap likely reflects a high false positive rate rather than precise TF assignment. In contrast, ReMap2022 annotations overlapped 38% of significant variants, reflecting lower genomic coverage but higher-confidence experimental support.

We tested whether specific TFs were enriched among significant variants. Using PWM-based predictions with local bit score filtering (see Methods, 32 TFs reached nominal enrichment (unadjusted P < 0.1; **Fig. 4**). These included broadly expressed architectural regulators such as NFYC, EBF1, and SREBF2, brain-enriched TFs such as RFX4, SCRT1, ETV1 and CUX2 (HPA brain-specificity ratio ≥ 2; see Methods), and a substantial number of C2H2 zinc-finger proteins (e.g., ZNF35, ZNF124, ZNF143, ZNF516, ZNF621), although no single TF family dominated the signal. Across the 32 TFs, the median number of significant variant overlaps was 5, and after BH correction, three TFs remained significant, NFYC, RFX4, and SCRT1 (FDR < 0.1, **Extended Data Fig. 7**). Applying the same test to ReMap2022 binding regions identified 22 TFs at nominal enrichment (P < 0.1), with KLF3, PHOX2B, RUVBL2 and SCRT1 showing the largest effect sizes (log2OR ≥ 3). Across these 22 TFs the median number of significant variant overlaps was 2.5 but no TF reached FDR < 0.1. Together, the PWM- and ReMap-based analyses indicate that allelic regulatory effects are distributed across many TFs rather than driven by a small number of dominant regulators.

**Figure 4:**
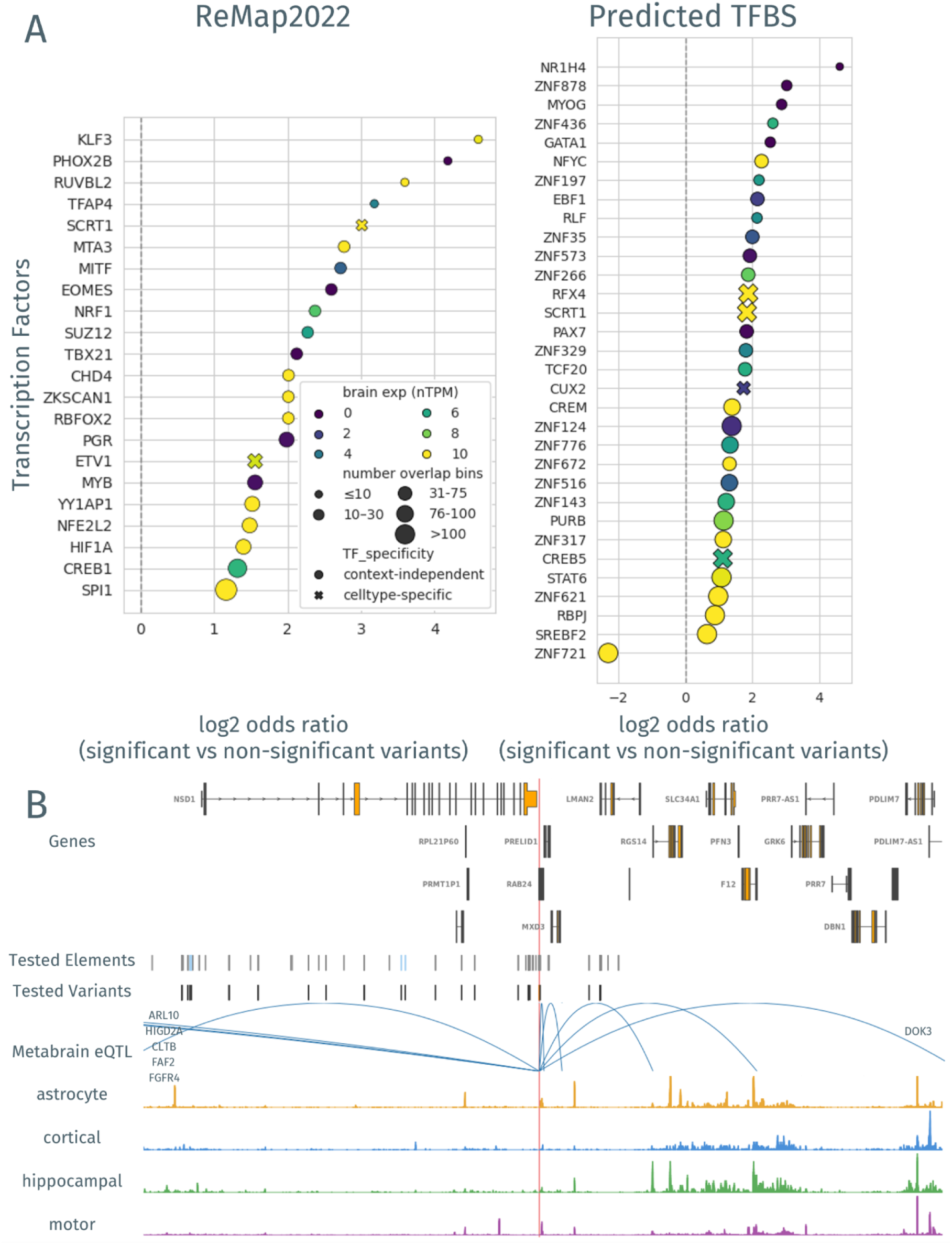
Activity driver and variant context. **a**, Effect-size based enrichment of TFBS in variants with significant effects in NGN2 derived neurons with experimental ChIP-Seq support from ReMap2022 (left) and predicted using FIMO and HOCOMOCO v13 (right). The color of the dots presents the expression in brain cell types defined by HPA. For visibility the range of shown expression is between 0-10 nTPM, so TF expression over 10 is presented as expression of 10. The “x” as shape is representing TFBS that have at least 1nTPM expression in brain and 2x the median expression (nTPM) in brain compared to the median in the other cell types according to HPA. The size of the dots reflects the number of variant overlapping TFBS found in all variants with readouts in NGN2 derived neurons and the y-axis reports the odds ratio of the enrichment. **b**, loci of rs1128287 (NC_000005.10:177301456:G:T) with the canonical isoform of the genes in proximity, the tested elements and variants, the overlapping MetaBrain eQTL from european cortex as arcs to the TSS of the associated gene, and ATAC-seq representation for astrocytes, cortical, hippocampal and motor neuron context.

For ReMap2022 annotations, precise motif positions and allele-specific binding consequences are not available as they are for PWM-based annotations. Therefore, we did not explicitly model gain versus loss of binding but treated overlap with experimentally supported binding regions as potential perturbation events. Although this simplification does not capture all possible binding changes, saturation mutagenesis studies suggest that sequence perturbations more frequently disrupt existing binding sites than create novel high-affinity sites ^75^, so under a loss-of-binding assumption an MPRA decrease in activity implies a TF that was activating, and an increase implies a TF that was reducing transcription. Across all ReMap TF-variant overlaps with at least two observations per TF, 41.5% of TFs were classified as context-dependent, 35.4% as predicted repressors, and 23.1% as predicted activators. Restricting to significant variant effects shifted the distribution towards repressors (48.8% predicted repressors, 30.2% context-dependent, 20.9% predicted activators). The equivalent PWM-based classification, which uses the sign of the bit-score delta together with the measured effect, yielded a much higher fraction of context-dependent TFs (70% across all overlaps; 23.2% in significant variants). Compared to motif-based predictions, ReMap-based annotations yielded fewer context-dependent TF classifications among all overlaps (41.5% vs. 70%), consistent with reduced annotation noise. Focusing only on significant allelic effects reduced the proportion of context-dependent classifications in both annotation strategies, indicating that variants with stronger regulatory effects tend to exhibit more consistent directional behavior.

To integrate the full set of annotations derived above with MPRA-derived effects, we examined the subset of 1,018 variants with measurable regulatory consequences in the MPRA assay. We note that 42% (424/1,018) had gnomAD AF > 0.01 (similar to proportion in the full dataset 47% (18,251/38,968)). Among significant variants, 355 variants were located within regions where either reference or alternative allele are annotated as dCREs, and 280 variants altered element activity sufficiently to switch between significant and non-significant regulatory states (dCRE reference only: 120, dCRE alternative only: 160). In addition, 213 variants overlap E2G annotations in brain-related cell types, and 23 variants overlap loci reported in the GWAS catalog, including 21 variants that both overlap a GWAS signal and a predicted TFBS. Together, these annotations placed a subset of MPRA-significant variants into regulatory and disease-associated contexts supported by independent lines of evidence (**Supplementary Table 5**).

We next highlight three representative examples that combine MPRA-detected regulatory effects with independent lines of annotation and phenotype evidence. An illustrative variant is rs17572795 (chr17:45,986,027 G>A; gnomAD AF = 0.14; log2FC = 0.95, ALT > REF), located within the ∼900 kb MAPT/KANSL1 inversion locus associated to Parkinson’s disease (PD) ^76^, which defines the common H1 (increased PD risk) and the minor H2 (protective) haplotypes ^77^. The alternative allele increased regulatory output in the MPRA with a predicted gain of ZNF83 motif affinity, although the element is not annotated as open chromatin in brain. In the PheWAS data, the alternative allele is protectively associated with Parkinson’s disease across three ancestry contexts (European β = −0.269, p = 1.4 × 10^-28 78^, South Asian β = −0.355, p = 5.8 × 10^-8^ 79, multi-ancestry β = −0.243, p = 4.5 × 10^-10 80^), matching the well-established protective role of H2 at one of the most consistently replicated PD loci ^77,81^. Because the inversion generates a single extended LD block, the variant appears as a haplotype-scale eQTL for MAPT, KANSL1, LRRC37A/A2, PLEKHM1, CRHR1, ARL17A/B, WNT3 and SPPL2C in MetaBrain ^82^ (e.g. LRRC37A2 β = 0.97 in cortex, p = 2 × 10^-261^, KANSL1 β = 0.82, p = 4 × 10^-164^), precluding single-gene causal assignment from cis-eQTL data alone. Current mechanistic candidates include KANSL1-mediated dysregulation of PD-relevant pathways via the NSL histone-acetylation complex ^83,84^ and LRRC37A/A2-driven astroglial inflammation ^85^. A non-brain-related example is rs45575136 in JAG1 (log2FC = 0.70; predicted gain of ZNF510 motif affinity), associated with reduced risk of Palmar fascial fibromatosis (Dupuytren’s disease; OR = 0.73, p = 7.2 × 10^-7^; UKB 500k WGS) ^86^ (**Extended Data Fig. 8**). Two orthogonal suggestive gene-assignment routes converge on two pathways associated to Dupuytren’s disease ^87^, Notch and Wnt. First, JAG1 itself as a canonical Notch ligand, a pathway activated across multiple organ fibroses ^88,89^. And MKKS as a nominal cis-eQTL in cultured fibroblasts (GTEx SuSiE, PIP = 0.0028, β = 0.26), encoding a chaperonin essential for basal-body and primary-cilium integrity whose disruption perturbs the ciliary WNT response ^90^, a well-established driver of fibroblast-to-myofibroblast transition in fibrotic tissue including palmar fascia ^91,92^. The protective direction of the effect and the convergent evidence from two independent gene-to-pathway routes make rs45575136 a plausible regulatory modifier of palmar fibroblast fate. In addition, we identified rs1128287 (gnomAD AF = 0.79, log2FC = -0.46), which is located in close proximity to MXD3 among other genes. Therefore it is providing a candidate mechanism for the repression of MXD3, which is detected by MetaBrain across cortical and subcortical brain regions (cortex EUR β = −0.554, p = 2.8 × 10^-65^), concordant with the MPRA result. MXD3 is a helix-loop-helix leucine zipper that is involved in Sonic Hedgehog-mediated GNP proliferation ^93,94^. The locus is associated with an increased serum urate level according to GWAS (β = 0.01, p ∼ 10^-13^) ^95^, attributed by Open Targets to RAB24, which encodes an atypical Rab GTPase. rs1128287 is in the credible set for an eQTL of that gene (dorsolateral prefrontal cortex, β = 0.890, p = 5.07×10^-44^, PIP = 0.322) ^96^. FOSL2 is predicted to gain binding affinity with the alternative allele (local bitscore 0.979), a brain-expressed AP-1 component associated with both, transcription enhancing ^97^ and repressing roles ^98^ in different contexts. NSD1, the adjacent Mendelian disease gene (Sotos syndrome; haploinsufficiency causes intellectual disability with ASD features in >83% of affected individuals ^99,100^, ends 1.2 kb upstream and shows only nominal MetaBrain brain eQTL effects (p ∼ 0.02), indicating that the common regulatory signal acts primarily on MXD3 and RAB24, not NSD1 (**Fig. 4b**). Although serum uric acid is significantly reduced in ASD children relative to typically developing controls, consistent with broader purine pathway dysregulation in ASD ^101^, a causal link between this common variant and ASD through either NSD1 or RAB24 remains speculative.

Overall, more than 19% of significant variants tested are located within dCREs in the cell type of interest, underscoring the relationship between regional regulatory activity and variant-level effects, but this does not imply that the remaining variants lack biological relevance. Of the 279 TFs whose PWM-predicted binding sites overlapped 663 significant variants outside dCREs, 97% are expressed in brain tissues (consensus nTPM ≥ 1; Human Protein Atlas), and 91% are broadly expressed across at least half of the 16 tissue groups examined. The expression breadth distribution of outside-dCRE TFs was indistinguishable from inside-dCRE TFs (Mann-Whitney p = 0.466) (**Supplementary Table 6**). Together, the near-ubiquitous tissue expression of these TFs supports the functional plausibility of MPRA-detected allelic effects even in regions without significant different activity compared to scrambled control sequences.

### Sequence-based model performance for predicting variant effects

To enrich functionally active rare variants within the design, we leveraged Enformer-predicted changes in chromatin accessibility (ΔDNase). From more than one million rare (gnomAD AF ≤ 1%) SNVs, we included variants with the highest activating effects, the highest repressive effects, and a random subset. This Enformer-guided selection strategy using mean absolute predicted change across all DNase-seq tracks resulted in a higher likelihood of identifying variants with significant allelic regulatory effects (odds ratio = 1.7), demonstrating that sequence-based predictions can serve as a heuristic for enriching functional variants, albeit with substantial false positives.

To evaluate how well current computational models predict allelic regulatory effects at single-nucleotide resolution and across the allele frequency spectra, we compared two state-of-the-art sequence-to-function models, Enformer ^102^ (200 kb sequence context; variant effects scored as mean and max absolute predicted change across all predicted targets as described in Methods), AlphaGenome ^14^ (1 Mb context; variant effects aggregated as mean and max absolute predicted differences across 11 genomic modalities and all predicted cell types as described in Methods), and NGN2 chromBPNet, a sequence-to-chromatin-accessibility model trained on ATAC-seq from the cell type used for this MPRA (see Methods). We included Combined Annotation-Dependent Depletion (CADD) score predictions ^102^ as a baseline. Further, it was shown previously that specialized models trained on MPRA data can substantially improve the predictive performance ^103^. Following this approach, we trained a model based on AlphaGenome’s fixed sequence encoder (see Methods) on our MPRA data using three-fold cross-validation. We assessed discrimination performance between significant and non-significant variants using ROC and PR analyses across all variants, variants in dCREs, allele frequency strata, activity quartiles, and NGN2 open chromatin regions (**Fig. 5**).

**Figure 5.**
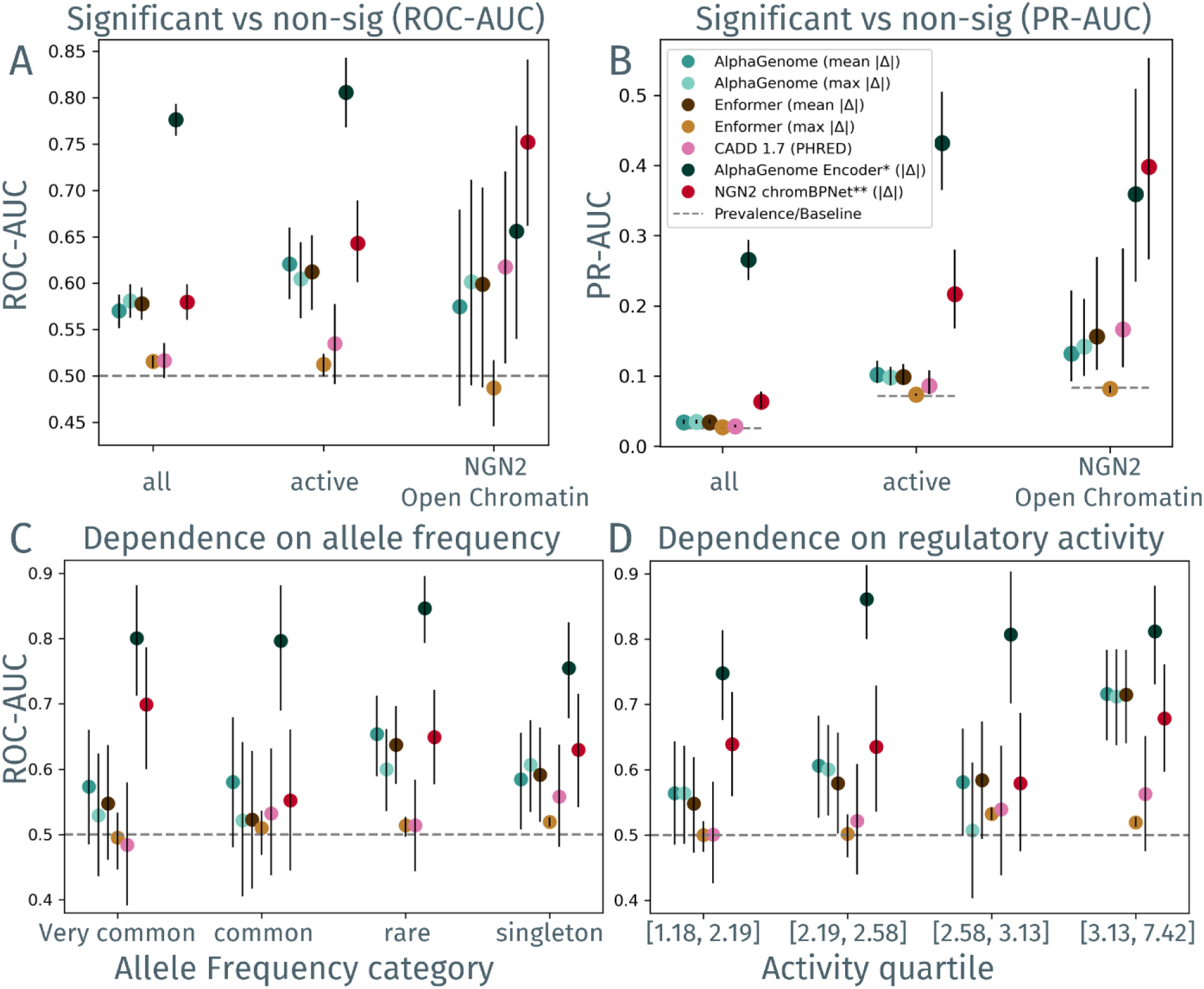
Sequence to function discrimination performance. **a**, Area under the receiver operating characteristics curve (ROC-AUC) and precision-recall curve (PR-AUC; **b**) for significant variant discrimination across three variant universes: all tested variants (all; n=38,968, pos=1,018), variants within dCREs with measured MPRA regulatory activity (active; n=2,711, pos=195), and variants within NGN2 open chromatin regions (NGN2 open chromatin; n = 453, pos=38). Models comprise AlphaGenome and Enformer (each evaluated using both mean and max absolute predicted variant effects), CADD 1.7, AlphaGenome Encoder*, and NGN2 chromBPNet**. Using variants in active regions **c**, ROC-AUC for model discrimination stratified by allele frequency and **d**, regulatory activity quartile of the enclosing element. Models marked * fine-tuned on MPRA data from this study; ** denotes a model pre-trained on cell-type-matched ATAC-seq from NGN2 excitatory neurons.

Using mean absolute predicted change as the variant effect score, Enformer and AlphaGenome exhibited modest
discrimination (ROC-AUC ∼0.57-0.58, PR-AUC ∼0.034), while CADD performed near random (ROC-AUC 0.52, PR-AUC 0.029), across all tested variants (n = 38,968; 1,018 positives) (**Fig. 5a**). NGN2 chromBPNet achieved similar ROC-AUC (0.58) but nearly double the PR-AUC (0.064), indicating better enrichment of true positives under class imbalance. Restricting the analysis to variants within dCREs (n = 2,711; 195 positives) improved performance for Enformer and AlphaGenome (ROC-AUC ∼0.61-0.62, PR-AUC ∼0.099-0.102) and chromBPNet (ROC-AUC 0.64, PR-AUC 0.22) (**Fig. 5b**).

Stratification of dCRE variants by allele frequency revealed substantial but inconsistent variation for Enformer and AlphaGenome (ROC-AUC 0.52-0.65; PR-AUC 0.053-0.147), with best discrimination for rare variants, while CADD showed no consistent pattern (ROC-AUC 0.48-0.56; PR-AUC 0.061-0.111). NGN2 chromBPNet achieved comparable ROC-AUC across strata (0.55-0.70) but consistently higher PR-AUC (0.13-0.28). The wide confidence intervals across all models limited interpretability (**Fig. 5c, Extended Data Fig. 8a**).

Stratification by regulatory activity of the enclosing element revealed an increase in ROC-AUC for Enformer and AlphaGenome for the highest activity quartile (ΔROC-AUC ∼0.15- 0.17 between lowest and highest quartile; Q1 ROC-AUC ∼0.55, PR-AUC ∼0.09-0.10; Q4 ROC-AUC ∼0.72, PR-AUC ∼0.17-0.18), indicating that sequence models more reliably identify allelic effects in strongly active regulatory contexts (**Fig. 5d, Extended Data Fig. 8b**). This trend was not present for CADD. NGN2 chromBPNet did not show the same activity-dependent ROC-AUC trend (0.58-0.68) but consistently ∼2-fold higher PR-AUC than Enformer and AlphaGenome across all quartiles (PR-AUC 0.17-0.29), suggesting cell-type-matched chromatin predictions enrich for true positives across activity levels rather than concentrating performance at the highest activity quartile (**Fig. 5d**). In NGN2 open chromatin regions, i.e. the native regulatory context of NGN2 chromBPNet, it achieved its highest point estimate (n = 453; 38 positives; ROC-AUC 0.75, PR-AUC 0.40), suggesting an advantage of cell-type-matched chromatin predictions in this specific context, though wide confidence intervals owing to the limited number of positive variants precluded stronger conclusions.

We note that the models compared here differ along two dimensions simultaneously: the biological readout on which they were trained (general epigenomic tracks for Enformer and pre-trained AlphaGenome; cell-type-matched chromatin accessibility for chromBPNet; MPRA activity directly for the fine-tuned model) and the degree of cell-type specificity (multi-cell-type for pre-trained models; NGN2-specific for chromBPNet and the fine-tuned model). These two dimensions of variation are not independently controlled in this comparison, and performance differences therefore cannot be attributed to any single factor. As expected given this alignment between training objective, cell type, and evaluation benchmark, the AlphaGenome encoder fine-tuned on our MPRA data consistently outperformed all pre-trained models across all stratifications (ROC-AUC 0.78 across all variants, 0.81 in active elements; PR-AUC 0.27 and 0.43, respectively) (**Fig. 5a, b**), with robust performance across allele frequency strata (ROC-AUC 0.76-0.85; PR-AUC 0.28-0.54) and activity quartiles (ROC-AUC 0.75-0.86; PR-AUC 0.36-0.52) (**Fig. 5c, d**). This result demonstrates that sequence representations encoded by AlphaGenome’s pre-trained encoder are sufficient to support MPRA-specific variant effect prediction when adapted through task-specific fine-tuning.

## Discussion

We performed a large-scale lentiMPRA to systematically characterize the regulatory landscape surrounding disease-associated genes in human NGN2-derived excitatory neurons, assaying over 27,000 candidate cis-regulatory elements and nearly 39,000 naturally occurring variants spanning the full allele-frequency spectrum. Our results reveal several interconnected findings about the architecture of neuronal regulatory variation, the relationship between AF and functional impact, the regulatory logic of transcription factor binding, and the utility of current computational models for predicting variant effects.

Consistent with emerging data from large-scale MPRA studies in neural contexts ^25,35^, we find that regulatory activity is sparse: only a minority of ENCODE SCREEN-annotated cCREs exhibited significant activity in NGN2-derived neurons, underscoring the inherent cell-type specificity of regulatory programs. Crucially, however, brain-annotated elements were enriched for active sequences, and, when active, displayed a near-balanced distribution of activating and repressive effects. Non-brain-annotated elements were instead strongly biased toward transcriptional repression in this context, a pattern consistent with chromatin-remodeling and co-repressor complex activity rather than active transcriptional enhancement. This context-dependent regulatory architecture mirrors findings from recent lentiMPRA studies in the developing cortex, which similarly highlighted strong cell-type and developmental specificity of enhancer activity ^25,104^. The substantial fraction of active elements lacking brain accessibility annotations in ENCODE SCREEN further underscores both the incompleteness of current epigenomic maps ^10,11^ and the capacity of MPRA to surface latent regulatory potential not captured by chromatin profiling alone ^26,105^.

A central motivation of this study was to examine how AF relates to regulatory functional impact. We identify significant allelic effects across the full allele-frequency spectrum, with comparable rates of detectable effects in common, rare, and singleton variants. While there is a modest monotonic decrease in the proportion of activating alleles from singletons to very common variants, absolute effect sizes show no strong frequency dependence. This is consistent with theoretical and empirical frameworks suggesting that regulatory variation is shaped by many alleles of modest effect distributed across the frequency spectrum ^17,106,107^. Importantly, the detectability of variant effects is more strongly governed by the baseline activity of the surrounding regulatory element than by AF per se. These observations have practical implications for variant prioritization: while AF remains a useful prior, local regulatory context provides substantially more information about the likelihood of a measurable functional consequence.

Roughly 90% of GWAS associations fall in non-coding regions and show modest enrichment within annotated cCREs, but this does not imply that most GWAS variants reside in cCREs ^15^. In our data, MPRA-significant variants were not enriched for overlap with GWAS catalog entries. GWAS reports lead SNPs in linkage disequilibrium, and fine-mapped credible sets typically span dozens of variants over tens of kilobases ^108^, so variant-level overlap is sparse by construction. Our design also sampled cCREs around disease-associated genes rather than GWAS loci, covering regulatory space that is only partly shared with common-variant architecture. The design is further restricted to a 50 kb window around annotated TSSs. This window captures most validated enhancer-gene connections ^109^, but misses long-range regulation, which can act over megabase distances ^110^. Variant-level overlap with the GWAS catalog is therefore an insensitive measure of regulatory relevance in this design.

Analyses of TF binding reveal a distributed and combinatorial regulatory architecture. Rather than a small set of dominant regulators, allelic effects are spread across numerous TFs, consistent with the billboard model of enhancer function ^111,112^ and recent evidence for heterotypic TF combinatorial binding in neural enhancers ^113^. TFBS enrichment and model-derived feature importance show only partial concordance, highlighting that motif presence alone is insufficient to predict regulatory output without accounting for binding cooperativity, chromatin state, and TF co-occupancy ^13,105,113–115^.

Sequence-based deep learning models (including Enformer and AlphaGenome) provide modest but consistent enrichment for variants with allelic MPRA effects, with performance improving markedly when restricted to elements with strong regulatory activity. An element activity-dependence would also be consistent with a mismatch between the genomic context used during model training as well as the out-of-context nature of MPRAs. This is supported by the improved performance of fine-tuning the AlphaGenome encoder on MPRA, demonstrating that pre-trained embeddings from large multi-task models encode sequence features that can be easily adapted to MPRA-derived signals using transfer learning. These comparisons can only be properly interpreted by acknowledging that different functional assays probe partially distinct aspects of regulatory mechanisms. MPRA quantifies the intrinsic transcriptional potential of isolated 270 bp sequence fragments at single-nucleotide resolution, but outside the endogenous chromatin environment and long-range regulatory architecture that shape in vivo allelic effects on e.g. chromatin accessibility or gene expression ^110^. No single functional readout constitutes a complete ground truth for regulatory variant effects, and models optimized for one assay are not expected to generalize to others without task-specific adaptation. Together, our data confirm that while sequence models and epigenomic annotations are valuable heuristics for prioritizing regulatory variants, large-scale functional assays remain essential for resolving fine-grained allelic effects in disease-relevant cellular systems and for training the next generation of context-aware predictive models.

Beyond their utility for benchmarking computational models, the significant allelic effects we identify across all allele-frequency classes, including among common and directly actionable variants for many individuals, represent a resource for mechanistic follow-up ^116–118^. In our dataset, a subset of significant variants overlaps fine-mapped brain eQTL loci, GWAS-associated signals, and E2G predictions in brain-relevant cell types, providing an immediately actionable list of candidates for in-depth investigation. More general, the functional catalog generated here offers a starting point for targeted CRISPR perturbation, allele-specific gene expression analyses, and TF binding studies at individual loci, translating allelic regulatory effects into mechanistic hypotheses of how disease-relevant genes are being modulated by standing variation.

## Methods

### MPRA design

The presented MPRA library was designed to investigate the cis-regulatory landscape around disease associated genes. As the library was supposed to be tested in neurons, high-confidence autism spectrum disorder (ASD) genes were selected from SFARI and literature-curated genes associated with neuropsychiatric disorders such as schizophrenia (n = 306, **Supplementary Table 1**) ^27^. To test how results from a single cell type extend to other cell types and diseases, we also collected a set of Congenital Heart Disease (CHD) genes (n = 164, **Supplementary Table 1**) from literature ^29–33^, a list of actionable clinical genes curated by the Center for Actionable Variant Analysis (CAVA; n = 48, **Supplementary Table 1**) and a set of randomly selected genes (n = 25, **Supplementary Table 1**). Genomic coordinates and Transcription Start Sites (TSSs) for all genes were obtained from GENCODE v42 ^119^. For each gene, we defined a 50 kb window around the union of all annotated TSSs and selected cCREs from the ENCODE SCREEN Registry (V3; excluding CTCF-only annotations) ^10^, excluding elements annotated as “CTCF-only”. cCRE sequences were oriented according to the gene strand. We selected 270bp from the center from the cCREs as design candidates. Control sequences, i.e. previously tested or otherwise prioritized sequences expected to show high and low activity respectively, as well as an IGVF oligos set for cross-comparison of assays (not within the scope of this study) were included (**Supplementary Table 6**).

Standing human genetic variations within these cCREs were obtained from gnomAD v3.1.2 ^66^. Variant discovery was restricted to SNVs overlapping each cCRE, excluding positions within 25 bp of the edge of each oligo. Across all selected cCREs design candidates, this yielded >1 million candidate variants, largely driven by rare alleles (gnomAD AF ≤ 1%). All common variants (gnomAD AF > 1%) were considered, while rare and singleton variants were prioritized as described below. Because of potential unwanted influences of the MPRA readout the chosen sequences were filtered, all sequences with an EcoRI or SbfI restriction site, a homopolymer longer than 10, UCSC simple repeat track (hg38/GRCh38) overlaps of more than 25%, a TSS overlap, or a CTCF overlap were excluded (https://github.com/kircherlab/MPRAOligoDesign). Elements and variants of the final library design are available as **Supplementary Table 6.**

### Enformer-based variant prioritization

Rare and singleton variants were prioritized separately for each gene group (neuro, cardiac, CAVA, random). For each group, 5,000 rare variants and 5,000 singleton variants were selected. Variant prioritization was performed using Enformer ^13^. For each variant, predictions were generated for both reference and alternative alleles in their native genomic context. Variants were ranked based on the maximum predicted allelic effect across DNase-seq tracks (maxΔDNase), following previously described scoring approaches ^13^. DNase-seq tracks were used due to their highest correlation with enhancer activity in MPRA assays.

From the ranked variants derived from disease genes, the top 3,500 variants, bottom 750 variants, and 750 randomly selected variants were chosen. For the variant set around random genes a smaller number, namely, 1,750 highest-scoring variants, 375 lowest-scoring variants, and 375 randomly selected variants were retained.

### Characterizing the composition of the dataset across gene groups

To assess whether the genes associated with the selected cCREs are expressed in the relevant cell type, RNA expression data were obtained from the Human Protein Atlas (HPA) Brain Atlas (version 25.0, Ensembl v109). For each gene, normalized transcript per million (nTPM) values for the cerebral cortex were extracted, and genes with ≥5 nTPM were classified as expressed. To annotate evolutionary constraint, cCREs were intersected with Regions of Constraint Conservation (RoCCs) derived from the Zoonomia project, downloaded from the UCSC goldenPath. A cCRE was classified as constrained if at least 1 bp overlapped a RoCC interval. For each gene, the proportion of associated cCREs overlapping constrained regions was computed. Interval operations were performed using pyranges (v0.1.4).

To contextualize the cCRE density and annotation properties of the MPRA gene sets, a genome-wide reference set was constructed. For all protein-coding genes in GENCODE v42 with available HPA cerebral cortex expression data, a 50 kb window around the union of annotated TSSs was defined and ENCODE SCREEN v3 cCREs (excluding CTCF-only annotations) were intersected, applying the same sequence quality filters used for the MPRA design (homopolymer, repeat, and TSS overlap exclusions). The resulting cCREs were annotated with brain and excitatory neuron SCREEN v4 biosample annotations, RoCC constraint, and linked gene expression as described above. This background set was compared against the neuro, cardiac, CAVA, and random gene sets across cCRE density, brain cCRE proportion, constraint, and gene expression metrics.

### Experimental procedures

#### Cell Culture

An engineered human induced pluripotent stem cell (hiPSC) line, WTC11, carrying a doxycycline-inducible neurogenin 2 (NGN2) transgene was used ^120^. WTC11 cells were maintained on matrigel-coated plates (Corning, 354277) in mTeSR1 medium (StemCell, 85850). Cells were routinely tested and confirmed negative for mycoplasma contamination. To initiate neuronal differentiation, doxycycline (final concentration: 2 µg/ml) was added to the differentiation medium to induce Ngn2 expression. The differentiation protocol has been previously described ^120^. Briefly, WTC11 cells were cultured in KnockOut DMEM/F-12-based pre-differentiation medium supplemented with doxycycline, N-2, BDNF, and NT3 for three days. This was followed by a switch to a Neurobasal-A and DMEM/F-12-based maturation medium for seven days, leading to the generation of post-mitotic excitatory neurons. On day 7, half of the medium was replaced with fresh maturation medium without doxycycline.

#### LentiMPRA Library Cloning and Sequence-Barcode Association

The lentiMPRA library construction is described in detail previously ^25,120^. The process essentially involved amplifying a synthesized oligo pool with a 5-cycle PCR and adding of minimal promoter (mP) and spacer sequences downstream of the cCRE. After purification, an additional 11-cycle PCR was performed to append a 15-nucleotide random barcode to each sequence. These amplified fragments were then ligated into the pLS-SceI lentiviral backbone vector using Gibson Assembly. To ensure comprehensive representation, approximately 16 million bacterial colonies were collected after transformation, aiming for an average of 200 unique barcodes per sequence. The association between each sequence and its corresponding barcode was subsequently determined by amplifying the sequence-mP-barcode fragments from the plasmid library and performing paired-end sequencing using a NextSeq Mid-Output 150PE kit with custom primers, generating roughly 123 million total reads.

#### Lentiviral Production, Infection, and Barcode Sequencing

Lentivirus particles, encoding the lentiMPRA library, were generated by transfecting the plasmid library into 293T cells utilizing the Lenti-Pac HIV expression packaging kit (GeneCopoeia, LT002). The crude viral solution was filtered through 0.45 µm PES unit (FB12566507, Fisher Scientific) and concentrated using Lenti-X concentrator (Takara Bio, 631232). The lentiMPRA library’s titration was performed on Ngn2-differentiated excitatory neurons to determine the optimal multiplicity of infection (MOI).

For each replicate, 25 million neurons were infected. Neurons were infected at an MOI of 100. To improve lentiviral infection in neurons, every 100 μl lentivirus was added with 6μl ViroMag (RL41000, OZ Biosciences). Neurons were infected on day 7 and continued to culture for another 7 days without medium change. An AllPrep DNA/RNA mini kit (Qiagen) was used for purifying DNA and RNA. Turbo DNase was utilized to remove contaminating DNA and unique molecular identifier (UMI) containing barcode-specific primer (P7-pLSmp-assUMI-gfp, from Gordon et al. 2020 ^121^) were used to reverse-transcribe the RNA with SuperScript IV (Invitrogen, 18090200). Genomic DNA and cDNA of each replicate were amplified and sample index and UMI were added using specific primers (P7-pLSmp-assUMI-gfp and P5-pLSmP-5bc-i#), with an initial 3-cycle followed by a 8-13-cycle PCR. Sequencing was performed with NextSeq High-Output SE75 or NovaSeqX 1.5B 100 cycles, and custom primers (R1, pLSmP-ass-seq-ind1; R2 (index read1 for UMI), pLSmP-UMI-seq; R3, pLSmP-bc-seq; R4 (index read2 for sample index), pLSmP-5bc-seqR2).

#### iPSC-derived neurons for ATAC-seq

NGN2-iNeurons were derived from WTC11 iPSCs containing a doxycycline-inducible NGN2 transgene integrated into the AAVS1 locus49. Cells were obtained from the laboratory of Dr. Yin Shen at the University of California, San Francisco (UCSF). iPSCs were maintained in StemFlex medium (ThermoFisher Scientific, A3349401) and underwent daily media changes. Cells were routinely passaged following dissociation with accutase (ThermoFisher Scientific, A11105-01) and replated on matrigel (Corning, 356231) coated plates. Media was supplemented with 10 µM ROCK inhibitor Y-27632 (Tocris, 1254) on the day of plating. For pre-differentiation, cells were replated on matrigel coated plates and cultured for 3 days in KnockOut DMEM/F−12 (ThermoFisher Scientific, 12660-012) medium supplemented with 2 µg/mL doxycycline (Sigma-Aldrich, D3447-500MG), 1x N-2 Supplement (ThermoFisher Scientific, 17502-048), 1x NEAA (ThermoFisher Scientific, 11140-050), 10 ng/mL NT-3 (PeproTech, 450-03), 10 ng/mL BDNF (PeproTech, 450-02), and 1 µg/mL mouse laminin (ThermoFisher Scientific, 23017-015). Cells underwent daily media changes during pre-differentiation and 10 µM ROCK inhibitor Y-27632 was supplemented on the first day of plating. Following 3 days of pre-differentiation, cells were dissociated with accutase and plated on Poly-L-Ornithine (Sigma-Aldrich, P3655-10MG) coated 10 cm plates. NGN2-iNeurons were cultured in BrainPhys neuronal media (StemCell Technologies, 5790) supplemented with 2 µg/mL doxycycline, 1X N-2 Supplement, 0.5X B-27 Supplement (ThermoFisher Scientific, 12587-010), 1X NEAA, 10 ng/mL NT-3, 10 ng/mL BDNF, and 1 µg/mL mouse laminin. A half-media change was conducted on day 7 without doxycycline.

#### ATAC-seq of NGN2-induced neurons

Assay for Transposase-Accessible Chromatin with high-throughput sequencing (ATAC-seq) was performed according to the ATAC-seq kit protocol (Active Motif, 53150). Approximately 100,000 NGN2-iNeurons were counted and pelleted by centrifuging at 500 x g for 5 minutes at 4°C followed by a wash in 100 uL of ice-cold DPBS (Thermo Fisher, 14190144) and an additional spin at 500 x g for 5 minutes at 4°C. Cells were lysed using 100 uL of ATAC Lysis Buffer (Active Motif, 53150) and centrifuged for at 500 x g for 10 minutes at 4°C. Tagmentation master mix was prepared using 25 uL Tagmentation Buffer, 2 uL 10x PBS, 0.5 uL 1.0% Digitonin, 0.5 uL 10% Tween 20, 12 uL H2O, and 10 uL Assembled Transposomes (Active Motif, 53150) and added to each sample followed by incubation on ice for 30 min. Tagmentation reaction was performed at 37°C for 1 hour shaking at 800 rpm on a benchtop heat block (Eppendorf Thermomixer C Model 5382). Following tagmentation, 250 uL DNA purification buffer (Active Motif, 53150) and 5 uL of 3M sodium acetate (Corning, 46-033-Cl) were added to the reaction mix, transferred to a DNA purification column, and centrifuged at 17,000 x g for 1 minute. DNA purification columns were washed with 750 uL of wash buffer (Active Motif, 53150) followed by elution in 35 uL of DNA purification elution buffer. Sequencing libraries were prepared using i7 and i5 index primers (Active Motif, 53150) and Q5 High-Fidelity 2x master mix (NEB, M0492) with the following PCR cycling conditions (72°C for 5 minutes, 98°C for 30 seconds, 11 cycles of 98°C x 10 seconds, 63°C x 30 seconds and 72°C x 1 minute, hold at 10°C). PCR reactions were purified with 1.2x SPRI select beads (Beckman Coulter, Cat. No. B23318). Libraries were ran on an Agilent 2100 Bioanalyzer using an Agilent High Sensitivity DNA Kit (Agilent Technologies, 5067-4626) to confirm average library size of 300 bp. Samples were sequenced at the UCSF Center for Advanced Technology (CAT) on an Illumina NovaSeq 6000 with an NovaSeq 6000 SP Reagent Kit v1.5 (100 cycles) kit (R1:50, R2:50, I1:8, I2:8) and standard Illumina sequencing primers.

#### ATAC-seq processing

ATAC-seq reads from NGN2-iNeurons were aligned to hg38 using the standard Encode Consortium ATAC-seq pipeline with default settings and pseudo replicate generation turned off (https://github.com/encode-dcc/atac-seq-pipeline).

#### Sensitivity and specificity of ENCODE SCREEN v4 tissue information

While sequences were originally designed from ENCODE SCREEN v3 (see above), for downstream annotation of cell-type and tissue specificity, we mapped the designed cCREs to SCREEN v4 annotations using the stable cCRE identifier shared between releases. This mapping preserved compatibility between v3 design coordinates and v4 biosample annotations. Among the tested cCREs, 721 (including 65 elements classified as significantly active in the MPRA) were no longer present in v4 and were excluded from cell-type-specific annotation analyses but retained for all other analyses.

To assign tissue-specific regulatory states, we downloaded the ENCODE SCREEN v4 “cCREs by Cell and Tissue Type” annotation files and extracted biosample-level cCRE annotations. We could not download cCRE information from SCREEN v4 for Fallopian tube, because of missing aggregation files. For each tissue, cCREs were classified as active if annotated as one of the following regulatory categories: CA-only, CA-H3K4me3, CA-TF, pELS, dELS, PLS. Binary indicators were generated per tissue type for each region to denote if the cCRE is considered active in this tissue.

#### Element activity and variant effect quantification

We used MPRAsnakeflow (v.0.4.5) with the default settings and BBMap ^122^ to assign the barcodes to sequences as well as counting these barcodes on RNA (reflecting expression) and DNA level (reflecting input) using the targeted DNA and RNA-Seq data ^123^. As quality filtering we used a minimal barcode threshold of at least 10 for oligos and both reference and alternative allele for variants. BCalm (v.0.9.0) was used for activity quantification on element level by performing a two-sided test for activity of an element being greater or smaller than a percentile threshold given by the activity distribution of the scrambled controls (mpra_treat (BCalm), 5^th^, 95^th^-percentile; FDR < 0.05), as well as statistically testing for the variant effects ^37^.

### Element-level analysis

#### TF enrichment in brain vs non-brain

To identify TFs differentially associated with MPRA-active elements that did versus did not overlap brain cCRE annotations, we annotated every tested cCRE with TF binding events derived from ReMap2022 ChIP-seq peaks ^39^. A TF was considered to overlap a cCRE if at least 10 bp of a ReMap peak overlapped the cCRE and the reported peak summit fell within the cCRE boundaries. For each cCRE and each TF, we recorded a binary indicator of whether at least one qualifying peak was present. To control for regions with many binding events and the GC content of the region we used them as covariates in the ridge-penalized logistic regression. Models were fit with scikit-learn’s LogisticRegression (L2 penalty, C = 1.0, lbfgs solver, max_iter = 3000). TFs with fewer than five positive cCREs across both groups combined were excluded to avoid unstable estimates. Confidence intervals and significance were estimated by non-parametric bootstrap (n=1000). Empirical p-values were corrected across TFs using the Benjamini-Hochberg procedure, and TFs with adjusted p < 0.05 were called as differentially associated.

#### TF enrichment in significantly active elements

As described above, we used ReMap2022 ChIP-seq peak annotations ^39^ to annotate the tested cCREs with TF binding events, and investigated the enrichment in regulatory elements exhibiting differential activity in the MPRA.

Significantly active elements were defined relative to scrambled controls (FDR < 0.05). For each TF, a 2x2 contingency table was constructed comparing the number of active and non-active cCREs containing at least one binding event for that TF. Enrichment or depletion was assessed using Fisher’s exact test (scipy.stats.fisher_exact), and p-values were adjusted using the Benjamini-Hochberg (BH) procedure. TFs with fewer than five total occurrences across all tested cCREs were excluded from statistical testing to avoid unstable estimates.

TFs were annotated with brain expression levels obtained from the Human Protein Atlas (HPA) ^124,125^. TFs with ≥5 nTPM in at least one brain region were considered expressed in brain tissue.

#### Feature modeling of element activity

To identify sequence and genomic features associated with MPRA element activity, we constructed a feature matrix at the cCRE level. ReMap2022 annotations were used to assign TF binding events to each cCRE using the overlap criteria described above. For each region, we computed (i) the total number of TF binding events and (ii) the number of unique TFs overlapping the region.

Additionally, we derived 7 binary features for brain and heart tissue from ENCODE SCREEN v4, indicating the cCRE group this region is associated to based on the epigenetic data available for this cell type, namely chromatin accessible only (CA-only), promoter like sequence (PLS), proximal enhancer like sequence (pELS), distal enhancer like sequence (dELS), CA and histone methylation (CA-H3K4me3), CA and transcription factor binding (CA-TF) and one column indicating if the region is considered to be overlapping any of the groups mentioned above.

Another binary feature is indicating an overlap with fine-mapped eQTL ^15^ data downloaded from the Finucane lab (https://www.finucanelab.org/data) as well as the phastCons score for each region fully covered by phastCons, the number of singleton variants per region to indicate population constraint and the distance of the region to the closest TSS (GENCODE v42).

All continuous features were standardized (z-score transformation) prior to modeling. To evaluate predictive performance while minimizing chromosomal information leakage, data were split using a chromosome hold-out strategy, with chromosomes chr16, chr18, chr20, and chr22 reserved as a test set (20% of chromosomes). An Elastic Net regression model (ElasticNetCV, scikit-learn) with 5-fold cross-validation was used to identify features associated with MPRA element activity. Similarly the performance was investigated with a balanced set of active and inactive sequences.

### Variant-level analysis

#### Variant coverage analysis

To assess the coverage of standing human genetic variation in the MPRA design, we reconstructed the set of variants that were present in gnomAD v3.1.2 within the tested cCREs. For each cCRE included in the final design, the valid variant window was defined by centering a 270 bp oligo on the SCREEN region and excluding 25 bp from each end, matching the edge exclusion applied during oligo design. Variants overlapping these windows were extracted from a local gnomAD v3.1.2 SQLite database and restricted to SNVs. Variants were stratified into four allele frequency classes: singletons (AC = 1), rare (gnomAD AF ≤ 1%), common (1% < gnomAD AF ≤ 5%), and very common (AF > 5%). The proportion of variants in each class that were included in the final MPRA design was computed to characterize the frequency spectrum of tested variation relative to the total available variation in the targeted regions. In total, we obtained effect measurements for approximately 83% of all gnomAD SNVs with AF >1% (n = 21,838 gnomAD v3.1.2) in the selected cCREs, including 10,036 variants with AF >5%. In contrast, for rare and singleton variants (AF ≤1%), we tested 20,560 variants, including 10,309 singletons, representing 1.7% of all naturally occurring rare variants in these cCREs (n = 1,188,427 gnomAD v3.1.2).

#### AF-effect modeling

Absolute variant effects (|log2FC| from BCalm) were modeled as a linear function of log_10_(gnomAD AF) by ordinary least squares (statsmodels OLS), with rare (AF ≤ 1%) and common (AF > 1%) variants fitted separately to account for the different sampling densities in the two regimes (**Extended Data Fig. 4**). Strand-pair and overlapping-cCRE duplicates (n = 284) were collapsed to one row per physical variant prior to fitting, retaining the orientation with the largest |log2FC|. As a robustness check, the same outcome was modeled across the full AF spectrum using a natural cubic spline on log_10_(gnomAD AF) with four interior knots (patsy cr() basis); the spline did not improve fit over the linear model in either the full dataset (F(3, 38679) = 2.24, p = 0.08) or the random subset (F(3, 21211) = 0.51, p = 0.68).

#### ReMap-based TF assignment to variant positions

To identify experimentally supported TF binding events overlapping tested variants, we used ReMap2022 ChIP-seq peak data. For each peak, the summit coordinate was extracted and extended by ±2 bp to generate a 5-bp window centered on the summit, representing the most probable TF binding location.

Variants were intersected with summit-centered windows using bedtools intersect (v2.31.1). A variant was considered overlapping a TF binding site if its genomic position intersected the extended summit window. For each overlapping event, the corresponding TF name was assigned based on the ReMap target annotation. No strand restriction was applied during intersection.

#### PWM-based TF prediction at variant positions

To identify TFBSs potentially affected by sequence variation, all MPRA oligo sequences were scanned using FIMO (MEME Suite) with HOCOMOCO v13 CORE motifs in MEME format ^126,127^ and motif occurrences as well as their genomic coordinates and strand orientation was used.

For each motif occurrence overlapping a tested variant, motif-sequence compatibility was quantified using the corresponding position weight matrix (PWM). To focus on sequence positions most likely influenced by the variant, we computed a localized PWM score within a 5-bp window centered on the variant position (±2 bp). If the variant was located near a motif boundary, the window was shifted to remain within the motif span. For both reference and alternative alleles, the observed PWM score within this window was calculated and normalized by the maximum possible PWM score for the same positions to obtain a relative local fit score. Strand orientation was accounted for by reverse-complementing sequences when necessary.

Motif occurrences were retained if the maximum relative local score across reference and alternative alleles was at least 0.75, ensuring strong local compatibility for at least one allele. The predicted effect of a variant on TF binding was quantified as the difference between alternative and reference local PWM scores. After filtering, an average of 5.8 motif occurrences per variant remained.

Motif identifiers were mapped to gene symbols using HOCOMOCO annotation metadata, and TFs were annotated with expression levels from the HPA brain RNA-seq dataset. For each variant, the TF with the highest expression in brain tissue among retained motif hits was selected as the representative candidate regulator. For statistical significance testing TFs with ≥ 5 overlapping variants were considered and Fisher’s exact test was used together with adjustment for multiple-testing using Benjamini-Hochberg.

#### Transition/transversion analysis

Single nucleotide variants were classified as transitions (A-G, C-T) or transversions (A-C, A-T, G-C, G-T) and stratified into four allele frequency categories based on gnomAD v3.1.2 annotations, singletons (AC = 1), rare (AF ≤ 1%), common (1% < AF ≤ 5%), and very common (AF > 5%). The proportion of transversions among the set of significant variants within each allele frequency category was computed and two-sided Fisher’s exact test was used to estimate odds ratios and p-values for each category.

#### Identifying expression of genes in brain

To prioritize biologically relevant TFs, expression levels in human brain tissue were obtained from the HPA, version 25.0 (Ensembl v109). RNA expression data were downloaded from the HPA Brain Atlas (“rna_brain_region_hpa.tsv”), which contains transcript expression levels summarized across 13 brain regions derived from RNA-seq analysis of samples from the Human Brain Tissue Bank.

For each gene, normalized transcripts per million (nTPM) values were aggregated across the following brain regions: amygdala, basal ganglia, cerebellum, cerebral cortex, choroid plexus, hippocampal formation, hypothalamus, medulla oblongata, midbrain, pons, spinal cord, thalamus, white matter. For three genes, namely *LINC01583*, *LINC01033*, *LINC01626* no expression was found.

For each TF, the maximum nTPM value across all brain regions was used to represent brain expression, thereby capturing region-specific high expression patterns. TFs with expression ≥5 nTPM in at least one brain region were considered expressed in brain.

#### Transcription factor tissue enrichment classification

To classify TFs into brain-enriched and non-brain enriched, we used RNA tissue consensus data from the HPA. Tissues were grouped into brain and non-brain categories. For each gene, we calculated the median nTPM across brain tissues and across all remaining tissues. A TF was classified as *celltype-specific* if (i) median brain expression was at least two-fold higher than median non-brain expression and (ii) median brain expression was ≥ 1 nTPM. All other TFs were classified as *context-independent*.

### Fine-tuning an AlphaGenome encoder model on MPRA data

#### Data preparation and cross-validation strategy

The training dataset comprised 65,308 sequences (270 bp) with associated activity measurements normalized to a scrambled sequence set. To reflect the experimental MPRA construct, sequences were extended by adding fixed adapter sequences (15 bp on each side), resulting in 300 bp sequences. During model input generation, an additional 32 bp promoter sequence matching the experimental design was appended, yielding a total input length of 332 bp per sequence. All sequences were one-hot encoded prior to model input. To prevent information leakage between training and test sets, sequences were grouped by genomic origin using chromosome-level grouping, ensuring that sequences derived from the same genomic context were assigned to the same fold. For each group, summary statistics were computed, including sequence count, mean GC content, mean regulatory activity, number of significantly active sequences (adjusted P value < 0.05), and number of variant-containing sequences.

Three cross-validation folds were generated using a greedy optimization procedure to balance multiple properties across folds. Specifically, folds were constructed to achieve comparable distributions of (i) total sequence counts, (ii) number of variant-containing sequences, (iii) number of active sequences, (iv) cumulative GC content, and (v) cumulative activity levels. Groups were iteratively assigned to folds in descending order of size and complexity, minimizing a global imbalance score defined as the weighted sum of squared deviations from target distributions across folds. This procedure resulted in three folds with closely matched distributions across all considered properties.

#### Model architecture

We used the pretrained AlphaGenome encoder as a sequence feature extractor and added a custom prediction head to model MPRA activity. The encoder processes the full input sequence and generates a low-resolution representation. These embeddings were layer-normalized, flattened across sequence positions, and passed through a multilayer perceptron (MLP) with a hidden size of 1,024, ReLU activation, and dropout (0.1). A final linear layer predicted a single continuous activity value per sequence.

#### Training procedure

For each cross-validation fold, one-fold was held out as test data, while the remaining two folds were split into training and validation sets (80/20). Models were trained using the Adam optimizer (learning rate = 1×10-5) with a batch size of 128 for up to 100 epochs. Mean squared error (MSE) between predicted and observed activity values was used as the primary loss function. Model performance was additionally monitored using the Pearson correlation coefficient. Early stopping was applied based on validation Pearson correlation with a patience of 3 epochs and a minimum improvement threshold of 0.005. To improve robustness, data augmentation was applied during training. Sequences were randomly reverse complemented with a probability of 0.5 to enforce strand invariance. In addition, sequences were randomly shifted by up to ±15 bp using circular shifts (wrap-around), introducing positional variability while preserving sequence composition. These augmentations were applied only to the training data.

#### Variant effect prediction and cross-validated scoring

For downstream analyses, the trained models were used to predict regulatory activity for both reference and alternative allele sequences. To ensure strict separation between training and evaluation, predictions for each sequence were obtained exclusively from the model for which that sequence belonged to the held-out test fold. Variant effect scores were computed as the absolute difference in predicted activity between reference and alternative alleles.

#### Variant effect prediction - AlphaGenome, Enformer, CADD

AlphaGenome is a sequence-based model published in 2025 which uses an input context size of up to 1,000,000 bases (1 megabase) and predicts multiple different experimental tracks for many cell types, like Enformer (200,000 bases context), the sequence-based model we used to prioritize rare and singleton variants (described above) ^13,14^. We computed variant effects across 11 genomic modalities for AlphaGenome: ATAC-seq, CAGE-seq, ChIP on histones, ChIP on transcription factors, DNase-seq, PRO-cap, RNA-seq, polyadenylation, splice junctions, splice sites, and splice site usage. For each modality, AlphaGenome’s recommended variant scorers compare predicted REF and ALT allele signal within assay-specific spatial masks, for instance, exons for RNA-seq, a 501 bp window centered on the variant for chromatin accessibility and TF binding, or the full gene body for splicing, restricting the comparison to the biologically relevant region. Scores are aggregated within each masked region using modality-appropriate statistics, such as log-fold changes in summed signal for expression and accessibility assays, or maximum absolute differences in splice site probabilities for splicing modalities. Since each modality is predicted across many cell lines and tissue types, cell-type-specific scores were summarized per feature using max, mean, and median absolute differences, and quantile-normalized; the mean across all features was used as the final per-variant AlphaGenome score.

We used a similar approach for Enformer predictions using all available targets (https://github.com/calico/basenji/blob/master/manuscripts/cross2020/targets_human.txt), but unlike for AlphaGenome no quantile score was given. To handle the different distributions of variant scores for different assays, we used the common and randomly selected variants of the dataset to estimate a background distribution excluding the 99th quantile to normalize the feature values and compute a less biased global mean.

Combined Annotation Dependent Depletion (CADD) v1.7 scores were obtained by submitting a VCF file containing all tested variants to the CADD web server ^102^. For each variant, the PHRED-scaled CADD score was used for downstream analyses. The published AlphaGenome encoder was fine-tuned as described above and variant effect scores were computed as the absolute difference in predicted activity between reference and alternative alleles (|Δ|).

#### Performance evaluation

Model performance was evaluated on two variant sets: (i) all variants with sufficient MPRA readout (n = 38,968; 1,018 positives) and (ii) the subset of variants located within dCREs (n = 2,711; 195 positives), defined as regions with significant regulatory activity compared to scrambled controls. Receiver operating characteristic (ROC) and precision-recall (PR) analyses was performed to discriminate between significant and non-significant MPRA allelic effects. ROC-AUC and PR-AUC were computed for each model and dataset. Confidence intervals (95%) were estimated using stratified bootstrap resampling (n=2000), preserving the ratio of positive and negative variants.

To assess performance across variant properties, we stratified variants within dCREs by allele frequency and regulatory activity. Allele-frequency categories were defined based on gnomAD annotations as very common (AF ≥ 0.05), common (0.01 ≤ AF < 0.05), rare (AF < 0.01, AC > 1), and singleton (AC = 1). Regulatory activity bins were defined by quartiles of normalized MPRA activity. All analyses were performed in Python using scikit-learn and custom scripts.

### Phenotypic Annotation of MPRA Variants

#### GWAS Catalogue Overlap Analysis

To assess whether MPRA-tested variants overlapped previously reported GWAS signals, we queried variants and their linkage disequilibrium (LD) proxies against the GWAS Catalogue ^128^. For each variant, an LD backbone and proxy variants were identified using genotype data from 30,000 randomly selected unrelated individuals of European ancestry from the UK Biobank cohort ^129^. LD proxies were computed using a window size of 3,000 kb and an LD threshold of r² ≥ 0.5 and lifted over to GRCh38.

The resulting variant set was then intersected with a filtered version of the GWAS Catalog (v1.0.2) ^68^. For each variant or proxy overlapping a GWAS signal, associated study metadata were retrieved, including the reported trait, mapped trait annotation, Experimental Factor Ontology (EFO) term, study identifier, and PubMed identifier of the corresponding publication.

#### Multi-Source Association Lookup for Prioritised Variants

For the 90 MPRA variants prioritized on the basis of significant regulatory effects within active chromatin regions, orthogonal gene-linking evidence, or proximity to a canonical TSS, a cross-database association lookup was performed to retrieve published GWAS associations at these specific positions. Unlike the GWAS Catalog analysis above, this lookup used exact variant matching without LD expansion, enabling direct retrieval of association statistics for the tested alleles themselves.

Overlapping variants in summary statistics from four publicly available online resources, namely the AstraZeneca PheWAS Portal ^86^, the HuGeAMP Knowledge Portal Network (https://www.kp4cd.org/), the OpenGWAS Portal ^130^, and the Open Targets Genetics Portal ^131^, as well as from GWAS summary statistics described earlier ^132–136^. Because these resources use different reference genome builds, input variants were lifted over from GRCh38 (hg38) to GRCh37 (hg19) using bcftools liftover (v1.22; chain file downloaded from UCSC goldenPath) ^137^. Associations were filtered at the genome-wide significance threshold of p < 5 × 10⁻⁸.

Results from all sources were harmonized into a unified table comprising the input variant (in hg38 chrom-pos-ref-alt, rsID, and SPDI notation, alongside its hg19-lifted representation), the phenotype or endpoint, association statistics (p-value, effect size, standard error, and odds ratio where available), trait type (quantitative or binary), ancestral background of the study population, sample size, and provenance information (PubMed ID, DOI, source URL, and query date). Duplicate entries arising from the same variant–phenotype association appearing across multiple sources were removed (deduplicated based on phenotype, ancestry, sample_size, pmid). For associations derived from the UKBB–FinnGen meta-analysis (https://public-metaresults-fg-ukbb.finngen.fi/) ^138,139^, cohort-specific effect sizes, standard errors, and p-values from the UK Biobank and FinnGen contributing studies were additionally retained alongside the meta-analyzed statistics.

#### ChromBPNet-based prediction of variant effects and transcription factor motif annotation

To interpret the regulatory effects of MPRA-tested variants in NGN2-induced neurons, we utilized ChromBPNet, a dilated convolutional neural network featuring automatic bias correction. We trained ChromBPNet models on bulk ATAC-seq data from WTC11 iPSC-derived NGN2-induced neurons to predict variant effects and identify overlapping TFBS.

We processed the bulk ATAC-seq data using the ENCODE ATAC-seq pipeline (https://github.com/ENCODE-DCC/atac-seq-pipeline). To train the ChromBPNet models, we utilized overlapping peaks across replicates and sampled GC-matched negative regions to serve as the background. For bias correction, we applied the bias model from the ChromBPNet repository pre-trained on the ENCSR868FGK dataset. Models were trained via five-fold cross-validation splitting across chromosomes using the ChromBPNet repository (https://github.com/kundajelab/chrombpnet). Specifically, the models used 2,114-bp genomic sequences as input to predict the total read counts and the profile shape for the central 1,000-bp regions.

We centered each variant, extracted the surrounding 2,114-bp genomic sequence, and used it as input for the ChromBPNet model. Predicted variant effects were defined as the log fold-change in predicted accessibility across the central 1,000-bp region between the reference and alternative alleles. To account for strand symmetry, predictions were generated for both the forward sequence and its reverse complement, with the final predicted effect calculated as the mean log fold-change across both orientations.

To compare model predictions with MPRA measurements, predicted variant effects were correlated with experimentally measured allelic effects from the MPRA. Because most tested variants were located outside regions of open chromatin in NGN2 neurons, analyses were focused on subsets of variants overlapping ATAC-seq peaks derived from NGN2 neurons, where the model is expected to capture sequence determinants of accessibility more accurately.

To identify sequence features contributing to predicted variant effects, model interpretation was conducted using DeepLIFT/DeepSHAP (https://github.com/kundajelab/shap) to derive base-pair-resolution contribution scores from the count head of ChromBPNet. These scores were calculated separately for sequences containing the reference and alternative alleles, then averaged across cross-validation folds to highlight changes in sequence feature importance caused by the variant. To annotate TFBS overlapping the variants, contribution scores generated for the peak regions were clustered using TF-MoDISco ^140^ to identify recurring sequence patterns. Individual motif instances overlapping variant positions were then detected using FiNeMo (https://github.com/kundajelab/Fi-NeMo) on the alleles exhibiting higher predicted accessibility. Subsequently, motif identities were assigned by comparing these discovered motifs against known TFBS databases (https://github.com/vierstralab/motif-clustering). These motif annotations represent sequence similarity to known motifs and may not reflect the exact TF active in NGN2-induced neurons.

#### Software and statistical environment

Computation has been performed on the HPC for Research cluster of the Berlin Institute of Health. All analyses were performed using Python (v3.11) and R (v4.4.1). Statistical modeling of MPRA activity and variant effects was conducted using the BCalm package (v.0.9.0), which is based on the limma framework. Statistical tests were performed two-sided if not stated specifically. Machine learning analyses were performed using scikit-learn (ElasticNetCV, 5-fold cross-validation, (version)). Multiple testing correction was performed using the Benjamini-Hochberg procedure. Genomic interval operations were performed using BEDTools (v2.31.1), pybedtools (v0.12.0).

## Data availability

MPRA datasets generated in this study have been submitted to the IGVF portal (http://data.igvf.org) under the following accession number: IGVFDS1419ZPHD. The ATAC-Seq used in this study performed in NGN2 derived excitatory neurons is available on request. The visualization of open-chromatin peaks is available at the Gene Expression Omnibus under the accession number GSE113480.

## Code availability

Analysis and visualization code are available at GitHub (https://github.com/kircherlab/80K-Analysis).

## Acknowledgements

This work has been supported by the National Human Genome Research Institute (NHGRI) Impact of Genomic Variation on Function (IGVF) consortium grant number UM1 HG011966 (J.S., N.A., M.K.) and Deutsche Forschungsgemeinschaft (DFG) project number 464313370 (M.K., M.S.). Computation was performed on the HPC for Research cluster of the Berlin Institute of Health, the HPC@Charité and the OMICS HPC cluster of the University of Lübeck. We thank current and previous members of the Kircher lab for helpful discussions and suggestions, and all members of the IGVF MPRA focus group for their valuable contributions to the MPRA pipeline and standardization.

## Author information

### Contributions

Conceptualization: KS, CD, PD, TC, JS, MS, NA and MK; Data curation: KS, CD, QL, ZC, MH, SR and MS; Formal analysis: KS, PD, ZC, MH and SR; Funding acquisition: JS, MS, NA and MK; Investigation: KS, CD, QL, ZC, MH and SR; Methodology: KS, CD, PD, TC, ZC, MH, SR, MS, NA and MK; Project administration: MS, NA and MK; Resources: AK, CL, MP, JS, MS, NA and MK; Software: KS, PD, TC, ZC, MH, SR and MS; Supervision: AK, CL, MP, JS, MS, NA and MK; Validation: KS, CD and QL; Visualization: KS; Writing - Original Draft: KS, CD, ZC, MS, NA and MK; Writing - Review & Editing: KS, CD, PD, TC, QL, ZC, MH, SR, AK, CL, MP, JS, MS, NA and MK;

## Extended Data

**Extended Data Figure 1:**
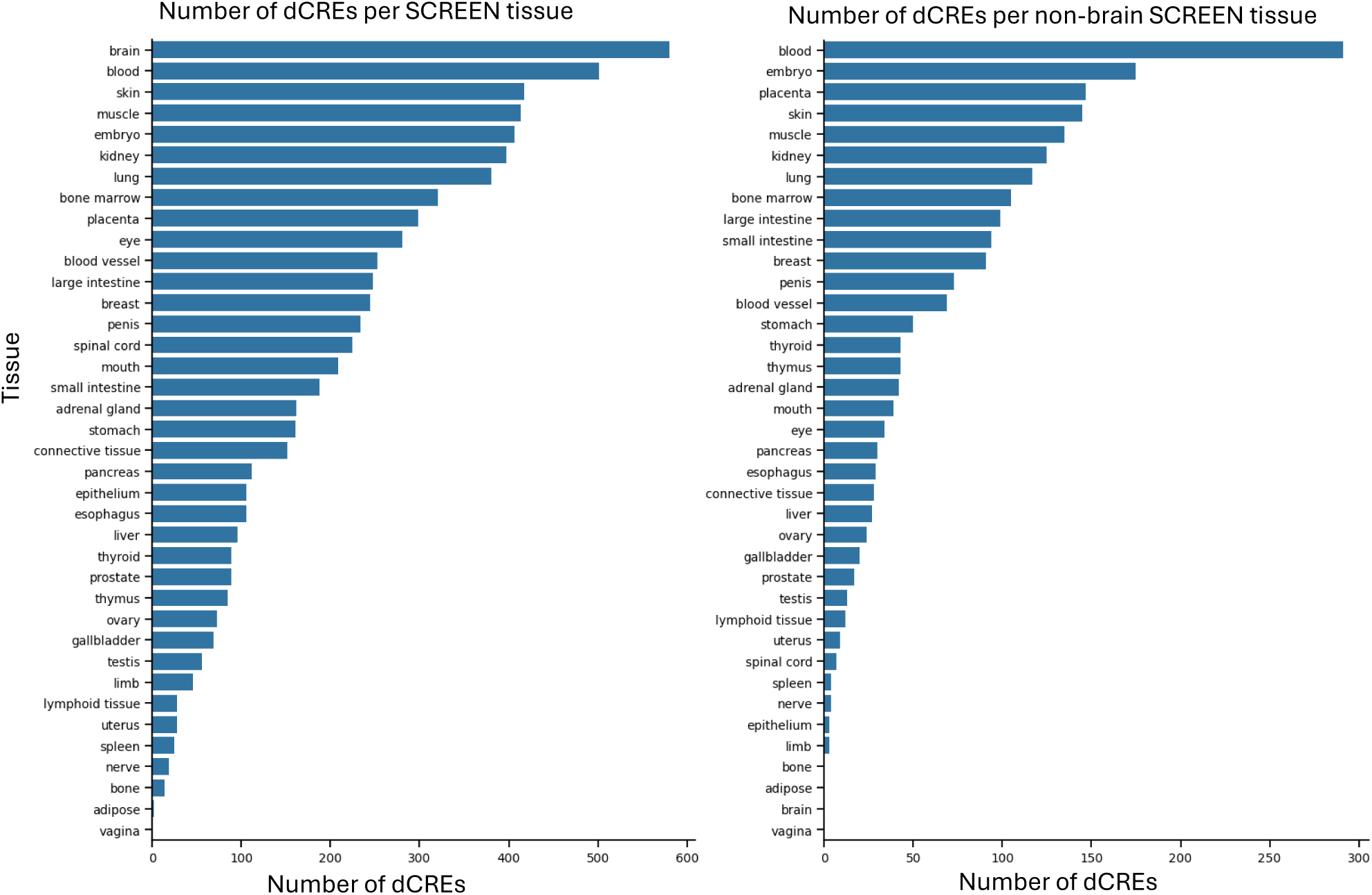
Number of dCREs per tissue annotation from SCREEN (left) and focusing on non-brain annotated dCREs (right).

**Extended Data Figure 2:**
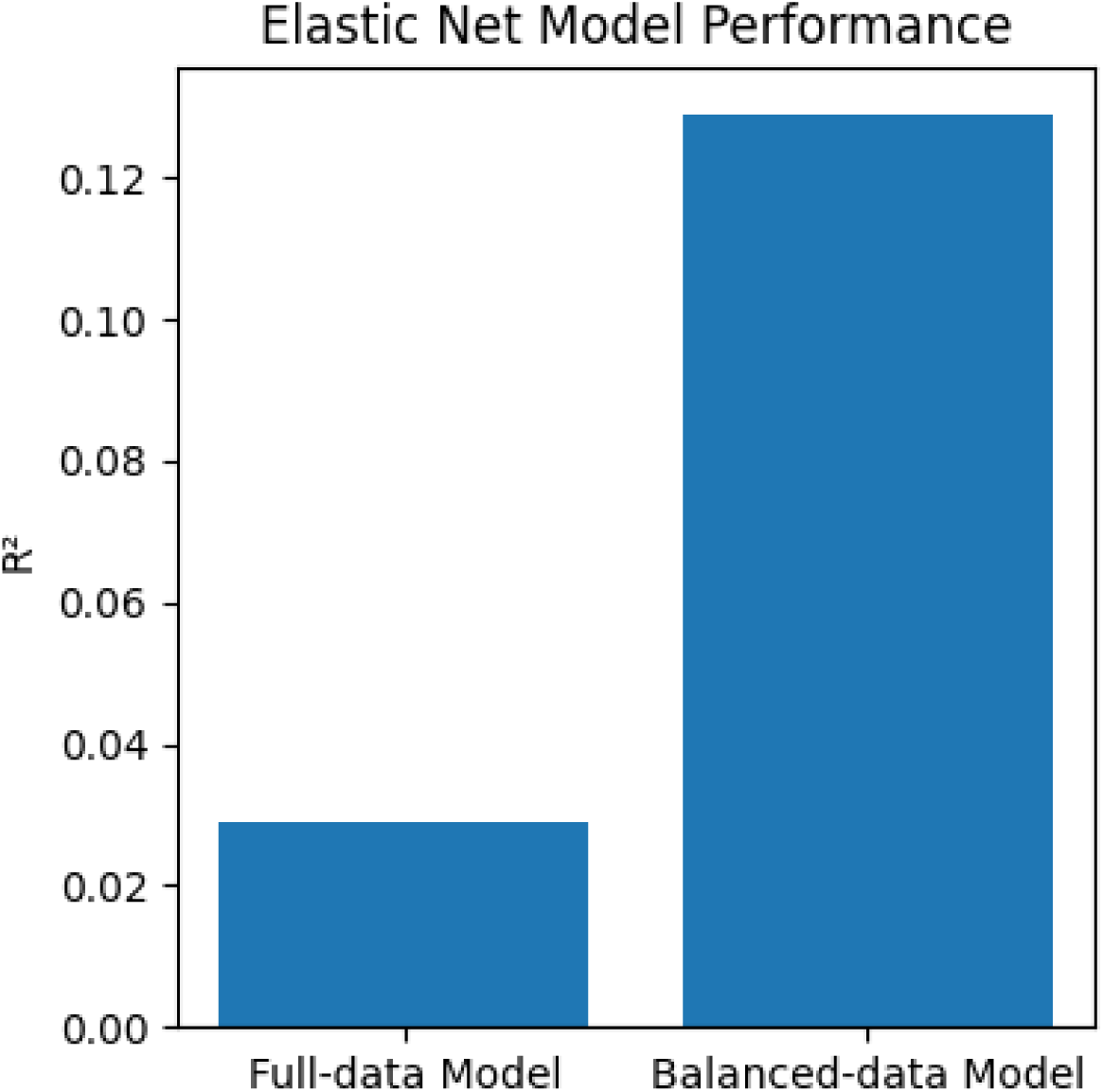
Elastic Net performance of the model using the full data set which is containing the biggest fraction of activity values around zero and a model trained on a subset of the data with equal amount of significant and non-significant cCREs.

**Extended Data Figure 3:**
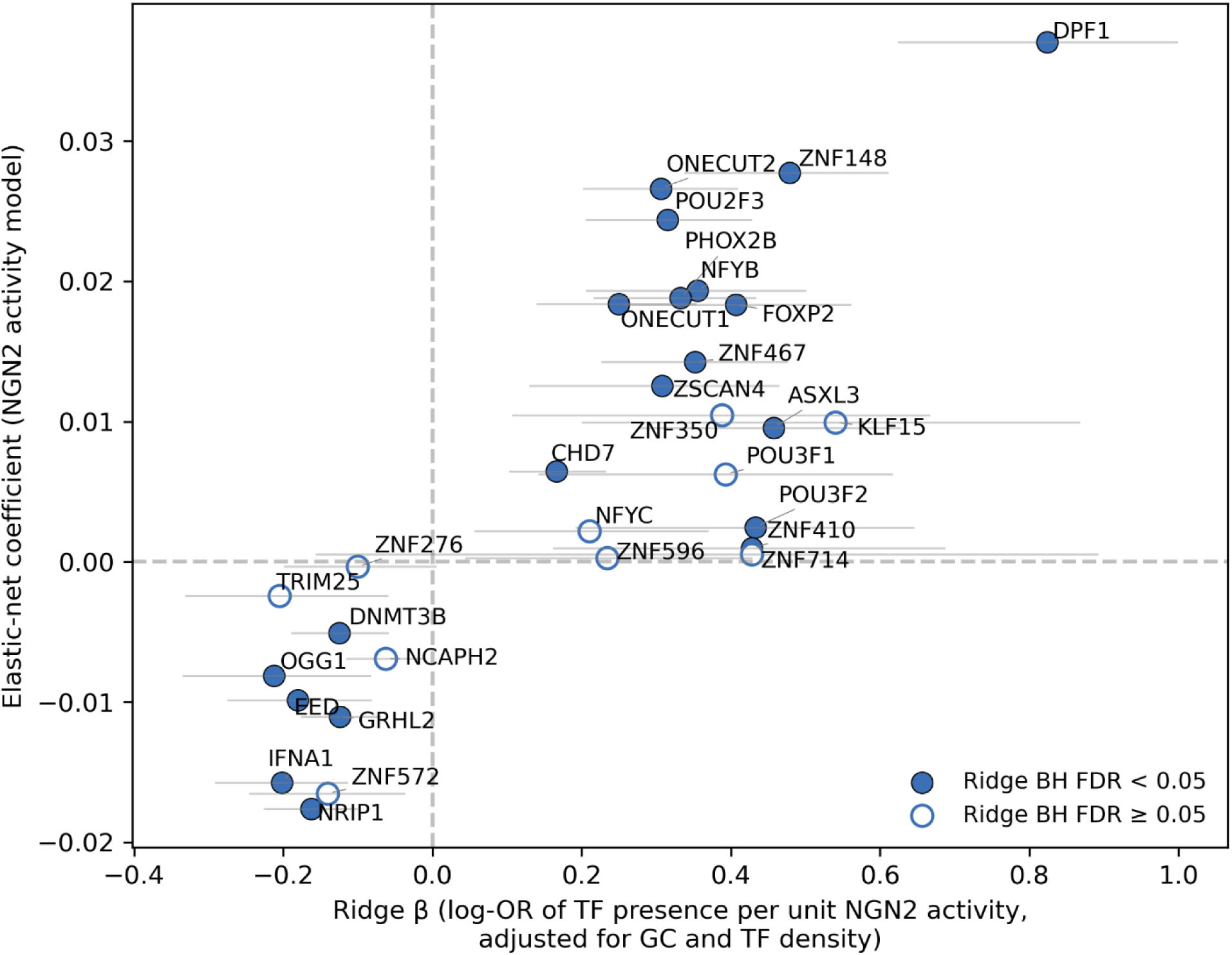
Each point is one of the 30 TFs that were both Fisher-significant in the dCRE vs non-significant comparison (BH FDR < 0.05; OR > 2 or < 0.8) and had a non-zero elastic-net coefficient. The elastic net coefficient and the ridge β from a per-TF logistic regression of TF binding presence on NGN2 normalized activity, fit on all 24,510 tested cCREs and adjusted for total TF density and GC content (TFBS_present ∼ activity + TF density + GC content); β is the log-odds of TF binding per unit normalized activity. Horizontal bars are 95% bootstrap CIs (1000 replicates). Filled markers reach bootstrap BH FDR < 0.05 across all tested TFs (n = 22/30) and open markers do not (n = 8/30).

**Extended Data Figure 4:**
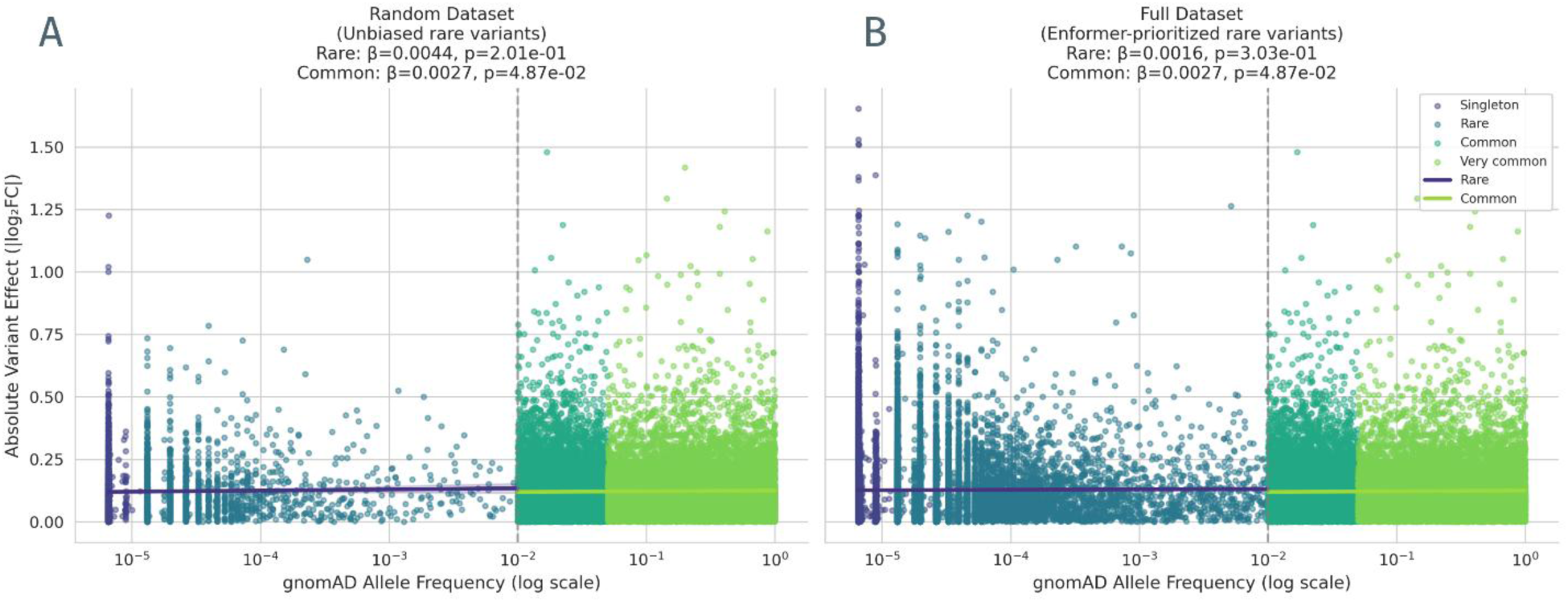
Absolute effect along log scaled allele frequency from gnomAD 3.1.2 for singleton (purple; AC=1), rare (blue;<=1%AF), common (dark green; >1%AF) and very common (light green;>5% AF) for the Enformer-prioritized rare variants (right) and the randomly selected rare variants (left).

**Extended Data Figure 5:**
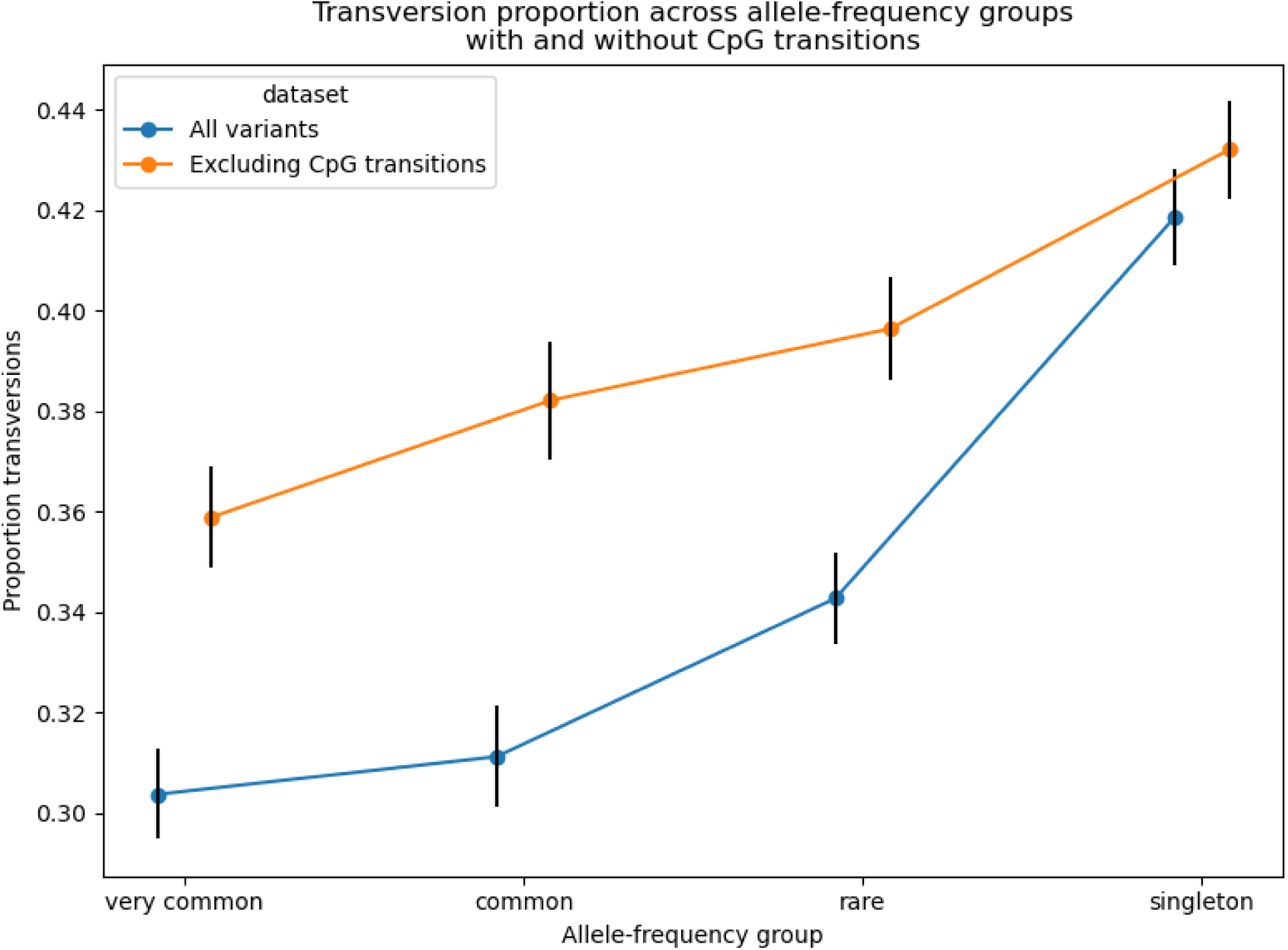
Proportion of transversion along the allele frequency (AF) groups very common (>5% AF), common (>1% AF), rare (<= 1% AF) and singleton (allele count = 1). Transversions show a significant enrichment with and without CpG associated variants.

**Extended Data Figure 6:**
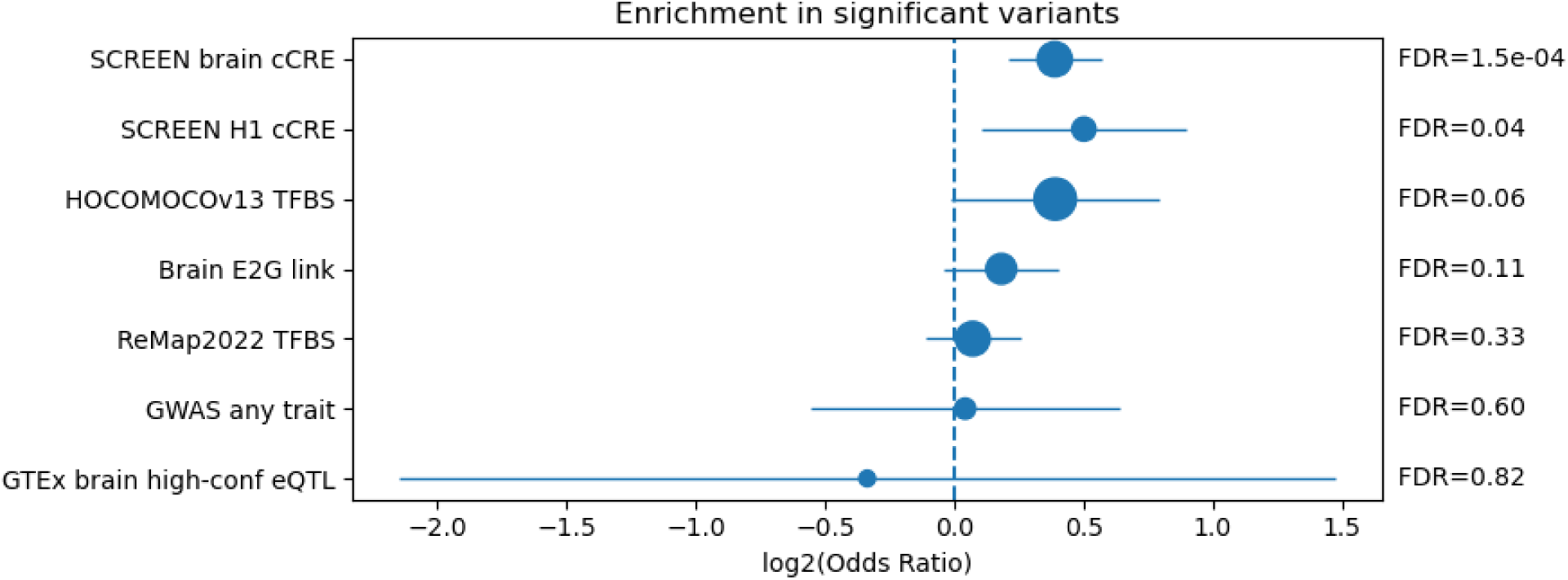
Enrichment of different annotation classes from SCREEN (brain, n=13,385, H1, n=1,566) (regulatory context), enhancer-to-gene (E2G) as gene linked (n=7,399), GWAS (n=873) or eQTL (genetic association) from fine-mapped eQTL data filtered using only brain tissue (n=119). Using Fisher’s exact test for odds ratios and estimating their confidence intervals and Benjamini-Hochberg for adjusting the p-value. The size is reflecting the number of variants within this class.

**Extended Data Figure 7:**
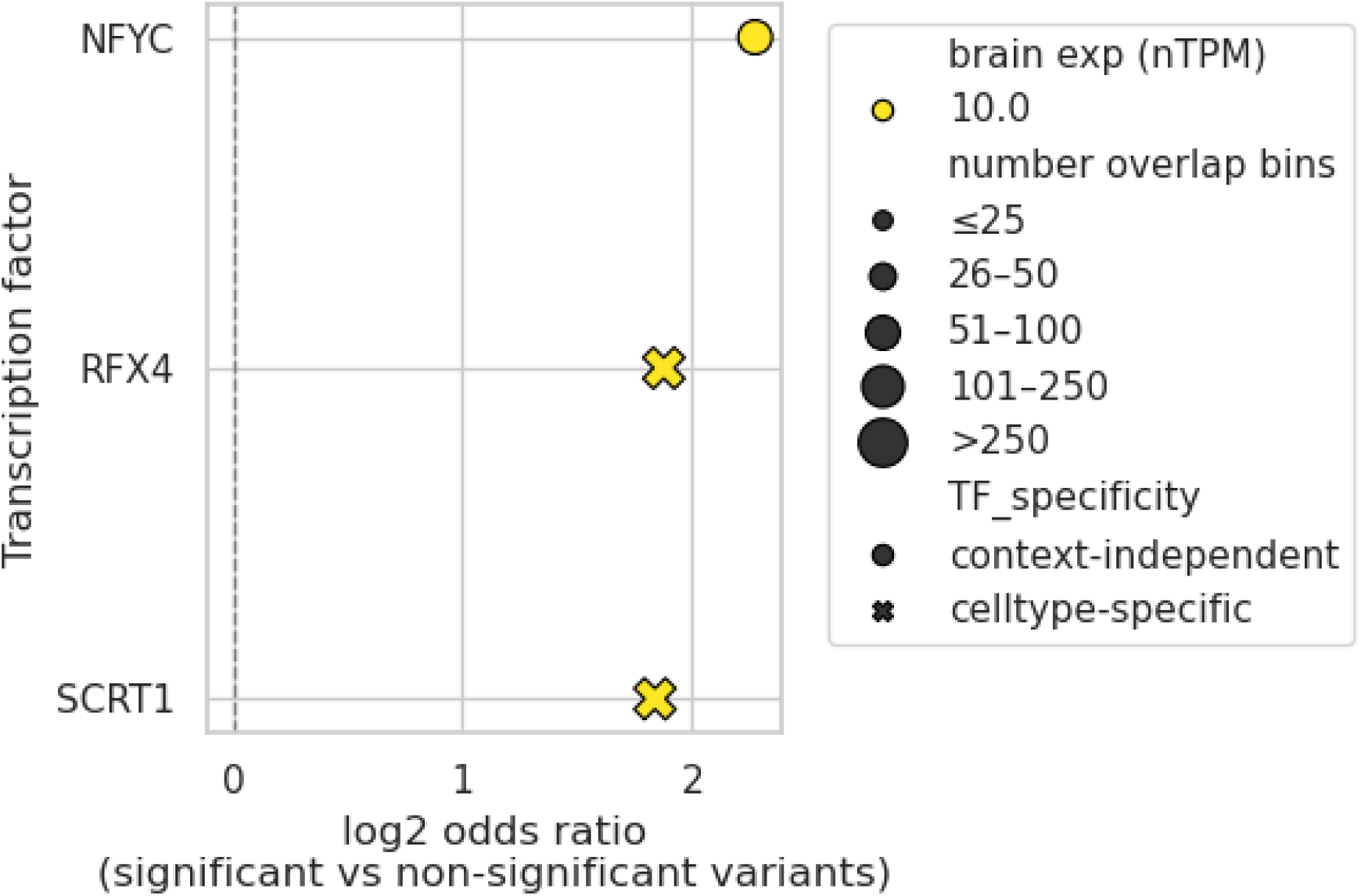
Enrichment of variant overlapping TFs from Hocomoco v13 using FIMO and bitscore based filtering for significant variants and adjusting for multiple testing with (bh adjusted FDR <0.1).

**Extended Data Figure 8.**
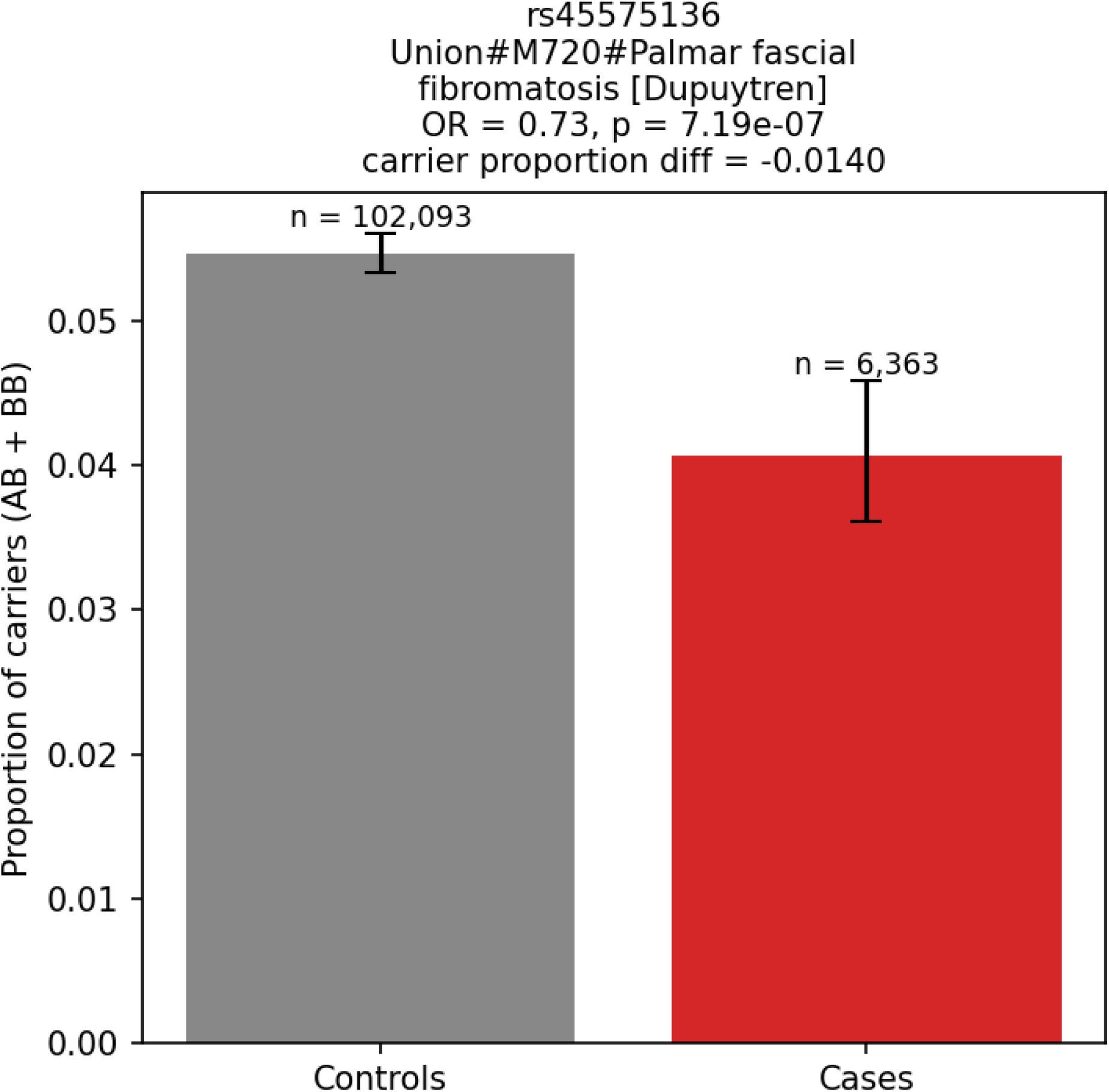
Proportion of individuals with Palmar fascial fibromatosis for both the individuals with the alternative allele both the control (gray; n=102,093) and in cases (red; n=6,363) derived from the AstraZeneca PheWAS Portal.

**Extended Data Figure 9.**
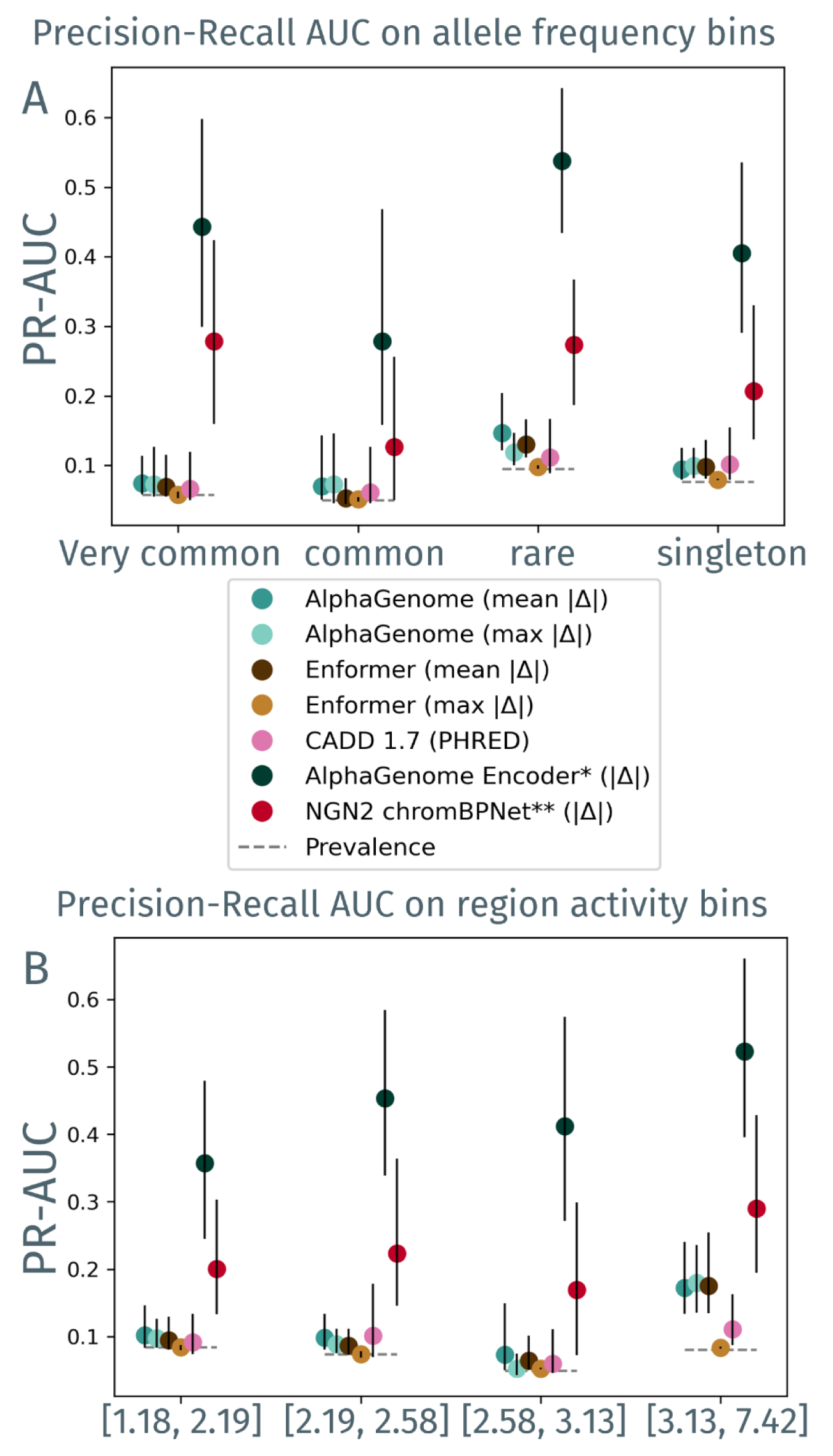
Sequence to function discrimination performance. Area under the precision-recall curve (PR-AUC) for significant variant discrimination of variants within dCREs with measured MPRA regulatory activity (active; n=2,711, pos=195) stratified by **a**, allele frequency and, **b,** activity of the associated region.

## Data

Supplementary Table 1: Gene with gene set group

Supplementary Table 2: Active brain vs active non-brain enriched TFs with ridge regression and bootstrap p-value

Supplementary Table 3: Elastic net model feature-coefficients

Supplementary Table 4: ReMap2022 enriched TFs sorted by odds ratio in active vs not active elements

Supplementary Table 5: Variant data with region annotations i.e. E2G, GWAS, eQTL

Supplementary Table 6: Metadata for elements and variants

Supplementary Table 7: Phenotypic annotation lookup of significant subset of MPRA variants

